# ActuAtor, a molecular tool for generating force in living cells: Controlled deformation of intracellular structures

**DOI:** 10.1101/2020.03.30.016360

**Authors:** Hideki Nakamura, Elmer Rho, Daqi Deng, Shiva Razavi, Hideaki T. Matsubayashi, Takanari Inoue

**Affiliations:** Johns Hopkins University School of Medicine, Department of Cell Biology and Center for Cell Dynamics, 855 N. Wolfe Street, Baltimore, MD, 21205, USA; Johns Hopkins University School of Medicine, Department of Biomedical Engineering, 855 N. Wolfe Street, Baltimore, MD, 21205, USA; Kyoto University Graduate School of Engineering, Department of Synthetic Chemistry and Biological Chemistry, Katsura Int’tech Center, Graduate School of Engineering, Kyoto University, Nishikyo-ku, Kyoto, 615-8530, Japan; Massachusetts Institute of Technology, Department of Biomedical Engineering, Cambridge, MA, 02139, USA

**Keywords:** Mechanobiology tool, mechanical force, organelle morphology, form and function, mitochondria, nucleus, non-membrane-bound organelles, membrane deformation, chemically inducible dimerization, optogenetics, synthetic biology, biomolecular condensates, intracellular mechanobiology

## Abstract

Mechanical force underlies fundamental cell functions such as division, migration and differentiation. While physical probes and devices revealed cellular mechano-responses, how force is translated inside cells to exert output functions remains largely unknown, due to the limited techniques to manipulate force intracellularly. By engineering an ActA protein, an actin nucleation promoting factor derived from *Listeria monocytogenes*, and implementing this in protein dimerization paradigms, we developed a molecular tool termed ActuAtor, with which actin polymerization can be triggered at intended subcellular locations to generate constrictive force in a rapidly inducible manner. The ActuAtor operation led to striking deformation of target intracellular structures including mitochondria, Golgi apparatus, nucleus, and non-membrane-bound RNA granules. Based on functional analysis before and after organelle deformation, we found the form-function relationship of mitochondria to be generally marginal. The modular design and genetically-encoded nature enable wide applications of ActuAtor for studies of intracellular mechanobiology processes.

## Introduction

Every phenomenon in nature is ruled by the mechanical laws of physics. Living organisms are no exception. Mechanical force regulates diverse biological processes including development (Mammoto et al., 2013), gene expression (Shivashankar, 2019), differentiation (Vining and Mooney, 2017), protein trafficking (Kassianidou et al., 2019), and dynamics of cellular and intracellular architecture including cell division (Itabashi et al., 2013; Elting et al., 2018), directed migration (Diz-Muñoz et al., 2013; Iskratsch et al., 2014) and organelle morphology regulation (Isermann and Lammerding, 2013; Helle et al., 2017). While cells sense and respond to extracellular physical forces mostly at the plasma membrane, recent studies implied active mechonoresponses in the intracellular milieu (Elosegui-Artola et al., 2017; Helle et al., 2017). Deficiencies or changes in cellular mechano-responses have been linked to diseased states including cancer (Ingber, 2003; Suresh, 2007; Isermann and Lammerding, 2013), collectively supporting the physiological importance of these mechanobiological processes. As such, there have been efforts to establish techniques for force manipulation at cellular and subcellular levels (Iskratsch et al., 2014; Liu, 2016; Norregaard et al., 2017). Among those techniques, the vast majority adopt one of two classes of probes for force exertion, i.e., physical or biological probes.

Physical probes include atomic force microscopy (Dufrêne et al., 2017), optical tweezers (Zhang and Liu, 2008), magnetic tweezers (Tanase et al., 2007; Sniadecki, 2010), pipette aspiration (Luo et al., 2013), micro- or nano-fabrication techniques such as patterned substrates (Nikkhah et al., 2012) and dielectrophoretic tweezers (Nadappuram et al., 2019). By being tractable and quantitative, these physical methods revealed a wide variety of cellular mechano-sensing mechanisms. However, there are limitations in introducing these probes into living cells (Norregaard et al., 2017), as well as in targeting a specific subcellular structure. While a few intracellular applications exist (Rai et al., 2013; Guet et al., 2014), their throughput is low, typically with a single location of force exertion at a time.

In contrast, biological probes such as motile microorganisms and engineered proteins can be introduced into cells with little effort. One example is the use of individual bacteria to apply force onto mitochondria (Helle et al., 2017). This study used pathogenic bacteria *Listeria monocytogenes* which enter host cells through phagocytosis followed by escape from the phagosome (Radoshevich and Cossart, 2018) (**Supplementary Figure 1a**). Once in the cytosol, the bacteria exploit host cell actin to move across the cytosol (Gouin et al., 2005), aiding them to infect neighboring cells for further spread across the host tissue. As a result, bacteria collide into subcellular structures by chance, thus applying force. While this technique realized relatively high throughput, it still lacks precise control over space and time of force exertion. Another example took advantage of inducible dimerization of two specific peptides that achieved connection between target organelle and microtubule motor proteins (van Bergeijk et al., 2015). Here, target organelles were transported along microtubules due to force generated by native microtubule motors, demonstrating maneuverability in both time and space. However, the force amount is apparently too small to induce visible deformation of target organelles, and force exertion is limited to where microtubules are. Heavy reliance on the native trafficking process thus limits its wide applications.

Networks of actin filaments generate force in cells and drive cellular functions such as endocytosis, cell migration and cell division. At the interface of biophysics and microbiology, the abovementioned bacterial strategy of generating force for motility has been well studied. To achieve this, the bacteria express ActA, a single-transmembrane protein, on the posterior half of their surface. ActA shares limited sequence similarity with two classes of proteins: an actin nucleator family, WASP, that functions as an Arp2/3-dependent actin nucleator (Bi and Zigmond, 1999; Skoble et al., 2000; Boujemaa-Paterski et al., 2001; Zalevsky et al., 2001), and actin effector family, vinculin/zyxin, which further facilitate efficient actin nucleation (Kocks et al., 1992; Golsteyn et al., 1997; Beckerle, 1998; Fradelizi et al., 2001). Concentrated ActA on the bacterial surface thus recruits the host actin machineries and triggers actin polymerization (Domann et al., 1992; Kocks et al., 1992, 1993; Pistor et al., 1994; Smith et al., 1995; Welch et al., 1997, 1998) (**Supplementary Figure 1b**). Resultant Arp2/3-dependent actin polymerization pushes the bacterial surface by the Brownian ratchet mechanism (Mogilner and Oster, 2003), to propel bacteria through the cytosol (Theriot et al., 1992; Cramer et al., 1994; Welch et al., 1997; Smith and Portnoy, 1997; Pantaloni et al., 2001) (**Supplementary Figure 1b**). *In vitro* reconstitution studies demonstrated that ActA is necessary and sufficient for the propulsion (Cameron et al., 1999; Loisel et al., 1999).

Due to increasing awareness of the physiological significance of intracellular mechanobiological events (Feng and Kornmann, 2018), there is a growing demand for a novel paradigm for force exertion onto intracellular objects with greater throughput as well as with higher spatio-temporal precision. In the face of this challenge, we rationally integrated an engineered ActA into induced dimerization techniques. Upon chemical or light stimulus, this molecular actuator rapidly deformed target intracellular structures as a result of *de novo* actin polymerization. We demonstrated striking deformation of both membrane-bound organelles and non-membrane-bound biomolecular condensates. This technique enabled assessment of the organelle’s morphology-function relationship, thereby demonstrating promise in probing mechano-responses inside living cells.

## Results

### Development of a molecular actuator for induction of actin polymerization and force generation

Inspired by the bacterial strategy of force generation through actin polymerization, we assumed that ectopic actin polymerization in cells results in *de novo* force generation (**Figure 1a**). Local concentration of actin nucleation promoting factors such as ActA is prerequisite for actin polymerization. Therefore, we sought ways to induce ActA accumulation in cells in a controlled manner. This could be achieved with chemically-inducible dimerization (CID) where chemical administration triggers protein-protein interaction (Komatsu et al., 2010; DeRose et al., 2013). More specifically, a soluble and minimally functional fragment of ActA is fused to one of the dimerizing proteins (e.g. FKBP), while the other half of the dimerizing pair (e.g. FRB) is fused to a signal sequence to localize the fusion protein to an intended subcellular location. Addition of a chemical dimerizer (e.g.. rapamycin) stimulates interaction between the two dimerizing proteins, which should result in the accumulation of the engineered ActA on the target membrane structure.

**Figure 1.**
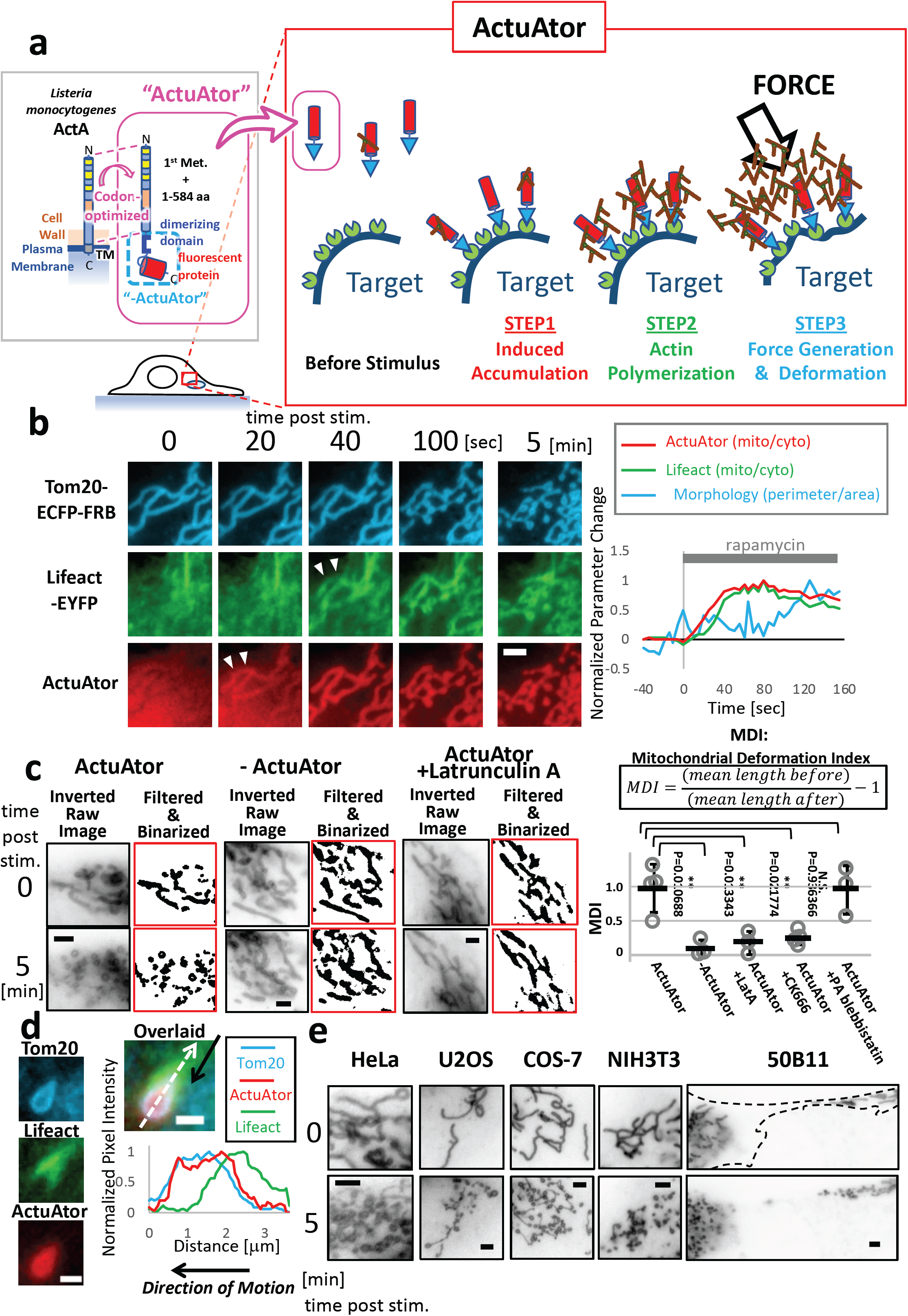
A novel molecular tool, ActuAtor, inducibly generates force inside living cells to deform mitochondria. a) Design scheme and steps of action of ActuAtor based on a bacterial protein ActA derived from *Listeria*. An engineered peptide, also referred to as ActuAtor, containing a domain from bacterial protein ActA along with a dimerizing domain, is expressed in the cytosol. The other peptide of inducible dimer is independently expressed on the surface of the target object inside the cell. The peptide is then inducibly accumulated onto the surface of the target object upon stimulus (STEP1), leading to actin polymerization (STEP2) that generates force onto the target (STEP3). These three steps mechanistically recapitulate the mechanism of actin-based propulsion of *Listeria* in host cell cytosol. b) ActuAtor deformed mitochondria upon stimulus. Left panel: Mitochondria deformation by ActuAtor in a U2OS cell is shown as sequential fluorescence images of Tom20-ECFP-FRB (mitochondria marker, cyan images), Lifeact-EYFP (polymerized actin marker, green images), and ActuAtor (ActA(1-584)-FKBP-mCherry, red images). Time after rapamycin addition is shown in seconds or minutes. First, ActuAtor peptide was inducibly accumulated on the surface of mitochondria 20 sec following rapamycin addition (white arrowheads in 20 sec panel). Actin polymerization was then detected in Lifeact channel (white arrowheads at 40 sec), followed by dramatic mitochondria deformation shown in cyan images (100 sec and 5 min). Scale bar: 2 μm. Right panel: Quantification of the data. Normalized ratio between ActuAtor signal intensity at mitochondria and in the cytosol (red), same normalized ratio of Lifeact signal (green), and normalized ratio between number of pixels at the perimeter and that contained in the entire area of binarized mitochondria image (cyan) are plotted against time after stimulus. Increase in the morphology parameter shown was preceded by the increase in ActuAtor signal and then that in Lifeact signal at the mitochondria. c) Mitochondria deformation by ActuAtor depends on actin polymerization by ActuAtor peptide. Left panel: Mitochondria morphology in HeLa cells before (upper panels, 0 min) and five minutes after rapamycin addition (lower panels, 5 min) are shown as inverted images (Inverted Raw Image), and as images binarized using top-hat morphology filter (Filtered & Binarized). Recruitment of the peptide lacking in ActuAtor peptide (-ActuAtor) and recruitment of ActuAtor in Latrunculin A-treated cells (ActuAtor +Latrunculin A) did not significantly deform mitochondria, while in intact cells, ActuAtor recruitment significantly deformed mitochondria into apparently fragmented morphology (ActuAtor). Scale bars: 2 μm. Right panel: Quantitative characterization of ActuAtor-induced mitochondria deformation. To describe the change in mitochondria morphology, mitochondrial deformation factor (MDI) was defined as change in average mitochondria length normalized by the length before stimulus quantified five minutes after stimulus (upper equation). MDI values were then used to characterize the deformation in different conditions (lower plots). Single data samples are plotted as grey circles, while mean ± standard deviation values are plotted as black markers. Compared to ActuAtor-induced deformation (ActuAtor, N=4 cells), lack in ActuAtor (-ActuAtor, N=3 cells), Latrunculin A pre-treatment (ActuAtor +LatA, N=3 cells), and Arp2/3 inhibitor, CK666 pre-treatment (ActuAtor +CK666, N=4 cells) significantly reduced the extent of deformation, while pre-treatment with myosin inhibitor, PA blebbistatin did not (ActuAtor +PA blebbistatin, N=3 cells). Statistical analysis was carried out by single-sided Welch’s *t*-test. **: P<0.05, N.S.: not significant. d) Deformed vesicular-shaped mitochondria move with trailing polymerized actin comet. Fluorescence images of a single moving vesicle are shown separately in the left panels. Overlaid image on the right and linescan plot along the white broken arrow indicates polymerized actin tail seen in green Lifeact channel is behind the vesicle. Scale bar: 1 μm. e) ActuAtor deforms mitochondria in similar manners in different cell types. Inverted fluorescence images of Tom20-ECFP-FRB right before (0 min) and five minutes after stimulus (5 min) are shown for different cell lines. Scale bars: 2 μm.

Toward this end, we first optimized codon usage of the extracellular domains (1-584 amino acids) of ActA (Kocks et al., 1992) to obtain efficient cytosolic expression with minimal side effects. Prior to this, we observed that expression of full length ActA in cells led to malformation of intracellular structures, and that the soluble fragment (1-584) in its original bacterial codon showed poor expression. We purified the codon-optimized, soluble ActA (1-584), and biochemically evaluated its actin nucleation efficiency. By performing a pyrene fluorescence assay, we could confirm robust actin polymerization with the modified ActA protein, which was comparable to a strong actin nucleator such as VCA domain of WASP (**Supplementary Figure 1c**). After confirming its potency as an actin nucleation factor, we generated a plasmid encoding the modified ActA fused to a dimerizing protein (FKBP) and a fluorescent protein (mCherry) (left panel, **Figure 1a**). When co-expressed with plasma membrane targeted FRB in cells, the ActA fusion protein (ActA(1-584)-mCherry-FKBP) started to accumulate at the plasma membrane upon addition of rapamycin, which subsequently led to actin polymerization and formation of actin-rich microspikes (**Supplementary Figure 1d**). A similar phenotype has been observed with constitutive anchoring of ActA to the plasma membrane (Friederich et al., 1995). We hereafter refer to the ActA-based molecular actuator as ActuAtor.

### ActuAtor induces mitochondria deformation that is independent of mitochondrial fission machinery

After validating the action of ActuAtor, we next applied ActuAtor to mitochondria for their deformation. To achieve this, we co-expressed ActuAtor with FRB fused to a mitochondria targeting peptide from Tom20. Upon ActuAtor accumulation in response to rapamycin addition, mitochondria morphology changed from tubular to apparently punctate, vesicularized shape within five minutes (**Figure 1b, Supplementary movie S1, S2**), which was not observed when FKBP without the ActA portion (-ActuAtor, left panel, **Figure 1a**) was instead accumulated at mitochondria (**Figure 1c**). We then evaluated the mechanism of deformation by observing the step-by-step molecular events. CID operation first led to ActuAtor accumulation, followed by the increased signal of Lifeact indicating actin polymerization, and then by the morphological deformation (**Figure 1b**). Accumulation of fluorescently-labeled Arp3 was also detected at the site of actin polymerization (**Supplementary Figure 2c**). In contrast, Arp2/3 inhibitor CK-666 treatment considerably weakened ActuAtor-induced Lifeact accumulation as well as deformation of mitochondria (**Supplementary Figure 2a**). Inhibiting actin polymerization by Latrunculin A pretreatment abolished the deformation (**Figure 1c**), while treatment with a myosin inhibitor, para-aminoblebbistatin (Várkuti et al., 2016), failed to perturb the observed deformation (**Figure 1c, Supplementary Figure 2b**). Together, these results supported the idea that ActA-based ActuAtor induced actin polymerization through Arp2/3, and the mitochondria deformation is caused by Arp2/3-mediated actin polymerization, but not through actomyosin contraction.

Actin polymerization has recently been implicated as a part of the physiological mitochondrial fission process (De Vos et al., 2005; Korobova et al., 2013; Manor et al., 2015; Li et al., 2015; Moore et al., 2016; Chakrabarti et al., 2018); polymerized actin is suggested to recruit Drp1, a GTPase that drives membrane constriction. Another study suggested that mitochondria respond to force applied to them by redistributing Mff, an adapter protein that recruits Drp1 to mitochondrial surface (Helle et al., 2017). We therefore investigated the requirement of endogenous Drp1 in ActuAtor-induced deformation of mitochondria by using a Drp1 KO mouse embryonic fibroblast. We found that ActuAtor deforms mitochondria even in the absence of Drp1, implying that the mechanism of this process is independent of endogenous mitochondria fission machinery (**Supplementary Figure 2d**).

We further evaluated off-target effects of the tool. ActuAtor induced intense actin polymerization at the site of action, suggesting potential unwanted effects on polymerized actin-based structures already existing in the cell. However, we found little effect of ActuAtor on pre-existing stress fibers, confirming few if any off-target effects on off-site actin-based structures (**Supplementary Figure 2e**). Other potential off-target effects are on organelles closely associated with mitochondria, especially endoplasmic reticulum (Rowland and Voeltz, 2012; Kornmann, 2013). By simultaneous imaging of endoplasmic reticulum and mitochondria in COS-7 cells, we found modest effects on network-shaped morphology of endoplasmic reticulum (**Supplementary Figure 2f**), while we cannot totally exclude potential local effects, including those on junctions between the two organelles.

### ActA offers the most effective deformation among other actin nucleators

It would be interesting to examine if and how ActuAtor probes consisting of actin nucleation promoting factors other than ActA differ in their performance of deforming mitochondria. We therefore generated four FKBP fusion proteins by replacing ActA with the following actin nucleators: full length N-WASP (N-WASP FL), constitutively active mutant of N-WASP (N-WASP ΔN) (Castellano et al., 1999), EspF_u_-5R (Sallee et al., 2008), and tandem SH3 domains of Nck (Nck SH3) (Taslimi et al., 2014). For each of the ActuAtor variants including the ActA-based one, we performed a cell-based CID assay, and quantified the degree of deformation (i.e., mitochondria deformation index, or MDI) by calculating the ratio of mitochondrial length before and after stimulus (**Supplementary Figure 3**). As a result, we observed some deformation for N-WASP ΔN (MDI of 0.54±0.56, indicated as mean±SD at 5-min post stimulation), and EspFu-5R (MDI of 0.43±0.14), while N-WASP FL and Nck SH3 had a marginal effect (0.15±0.31 and 0.13±0.13, respectively). With the MDI score of 1.28±0.40, ActA-based ActuAtor significantly outperformed the other nucleators. These results demonstrated that our strategy of concentrating ActA using the CID technique is generalizable to other actin nucleators, and that ActA was the most effective actuator among those tested. Thus, we used ActA-based ActuAtor for all remaining experiments.

While performing live-cell visualization of the ActuAtor-induced mitochondria deformation, we noticed that mitochondria often exhibited directed motion, and that these mitochondria associated with F-actin at their rear ends (**Figure 1d**), which is reminiscent of actin comets observed at the trailing edge of propelling *Listeria* (**Supplementary Figure 1b**). Of note, this asymmetric F-actin assembly did not colocalize well with ActuAtor which showed rather uniform distribution along the mitochondria. The distinct localization of ActuAtor and its downstream product, F-actin, implies the presence of a non-linear signal process at the level of Arp2/3-mediated actin polymerization, resulting in the symmetry breaking (where a spatially uniform input results in a polarized output). We speculate that the resultant polarized F-actin is a cause of the observed mitochondria motion, in a similar manner to the case of bacterial propulsion in host cells.

We next examined if ActuAtor-induced mitochondria deformation can be achieved in different cell types. In all five cell types we tested (HeLa epithelial cells, U2OS epithelial cells, COS-7 fibroblast-like cells, NIH3T3 fibroblasts, differentiated 50B11 neurons), efficient mitochondria deformation was observed (**Figure 1e**), demonstrating general applicability of ActuAtor.

### Optogenetic ActuAtor deforms mitochondria locally, reversibly, and repetitively

While CID allows for multicolor fluorescence imaging with ease, most CID tools including the rapamycin version we used here are practically irreversible in action. Also, chemical administration often lacks spatial control at a subcellular level. To gain more precise control of the *ActuAtor* operation, we next adapted light-inducible dimerization (LID) (Niu et al., 2016; Spiltoir and Tucker, 2019) for construction of optogenetic ActuAtor probes (opto-ActuAtor). In particular, we employed a light sensitive peptide, iLID, and its binding partner, SspB, which were fused to the codon-optimized ActA (1-584) and a mitochondria targeting sequence from Tom20, receptively. Upon blue light illumination to a sub-region of a target cell, opto-ActuAtor accumulation, actin polymerization, and mitochondria deformation were exclusively observed in the illuminated region (**Figure 2a**). When the stimulus light was turned off, mitochondria stopped deforming, and restored their initial tubular morphology (**Figure 2b**). The whole reversible process can be repeated by a second round of light on and off at the same region (**Figure 2b**). Together, opto-ActuAtor demonstrated reversibility, repeatability, and high spatial precision.

**Figure 2.**
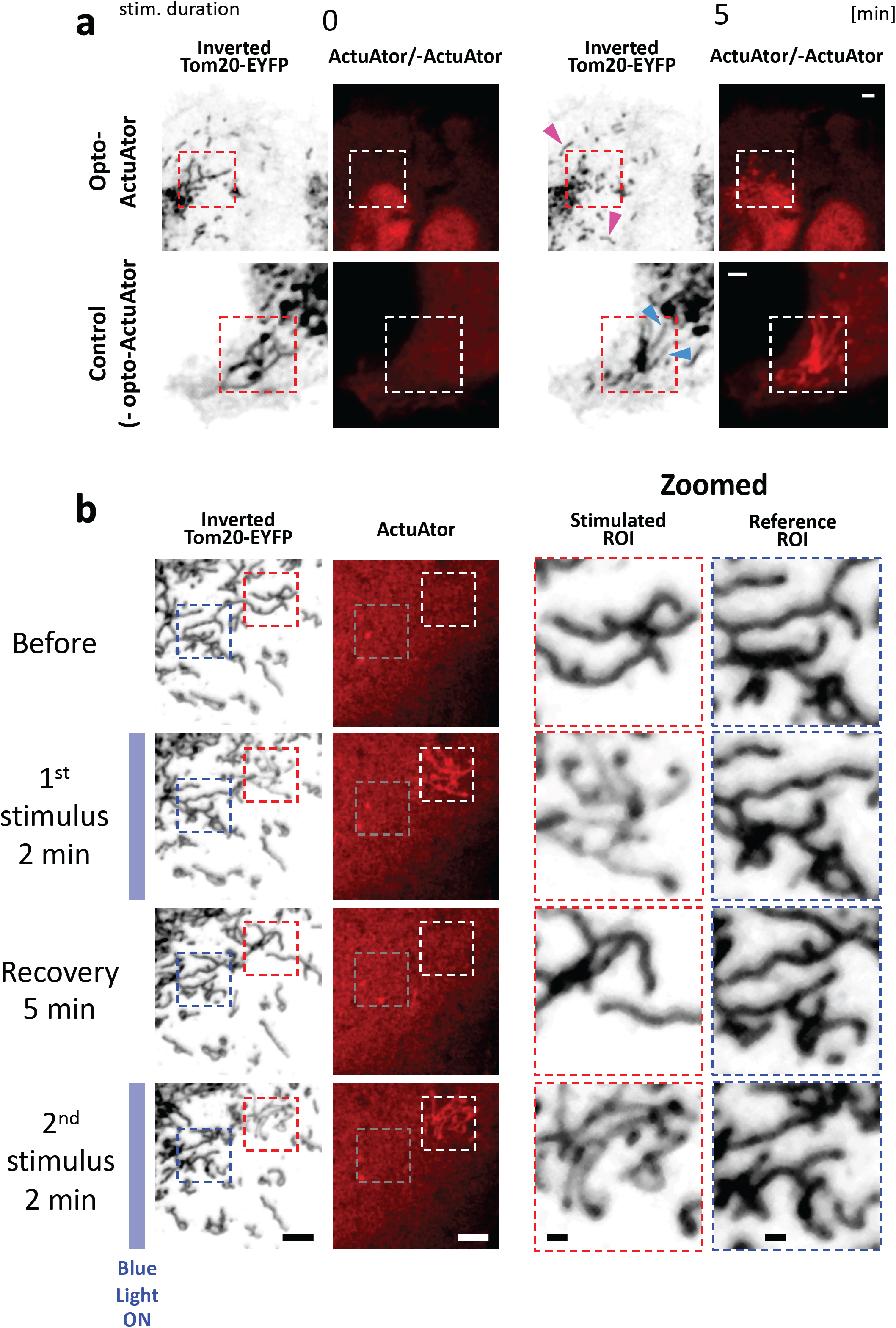
Light-inducible ActuAtor yields improved spatio-temporal control of mitochondria deformation. a) Light-inducible ActuAtor enabled local deformation of mitochondria. ActuAtor peptide was inducibly accumulated at mitochondrial surface by blue light-dependent dimerization between SspB and iLID in a U2OS cell. The light was irradiated in the square region shown by broken lines in the panels. Light was irradiated immediately before each acquisition during the stimulus. Scale bars: 2 μm. b) Repetitive and local deformation of mitochondria in U2OS cells by light. The same ROI (red broken-line square) was repeatedly stimulated. Mitochondria in stimulated ROI was deformed repetitively (Stimulated ROI in Zoomed panels, second and fourth panels from top) while no significant morphology change was observed in reference ROI (blue broken-line square, Reference ROI in Zoomed panels). Timing of the first and second stimulus is shown by blue bars on the left. Scale bars: 2 μm in left panels, 1 μm in Zoomed panels.

### ActuAtor induces deformation of Golgi apparatus

We further evaluated ActuAtor by directing the tool to another organelle, Golgi apparatus, using FRB fused to a fluorescent protein (ECFP) and to a Golgi-targeting sequence from Giantin (Komatsu et al. 2010). Rapamycin addition to cells co-expressing ActuAtor and ECFP-FRB-giantin triggered accumulation of ActuAtor at Golgi, and subsequent morphological changes of Golgi (**Figure 3a, Supplementary movie S3**). Fluorescence images of Golgi over time indicated a loss of dense spots within the Golgi cluster, which was quantified based on a linescan plot of the fluorescence intensity profile of this organelle (**Figure 3b**). These images were then processed by a top-hat morphology filter to capture subtle phenotypes; we were able to observe small particles emanating from the Golgi cluster (**Figure 3a**,**c**). When we performed visualization of Golgi morphology, F-actin, and ActuAtor, we could capture actin comet-like structures associated with the Golgi particles that were moving across the cytosol (**Figure 3d**). A subsequent linescan analysis of individual fluorescence signals confirmed that the actin comet-like structures were formed at the rear end of the Golgi particles.

**Figure 3.**
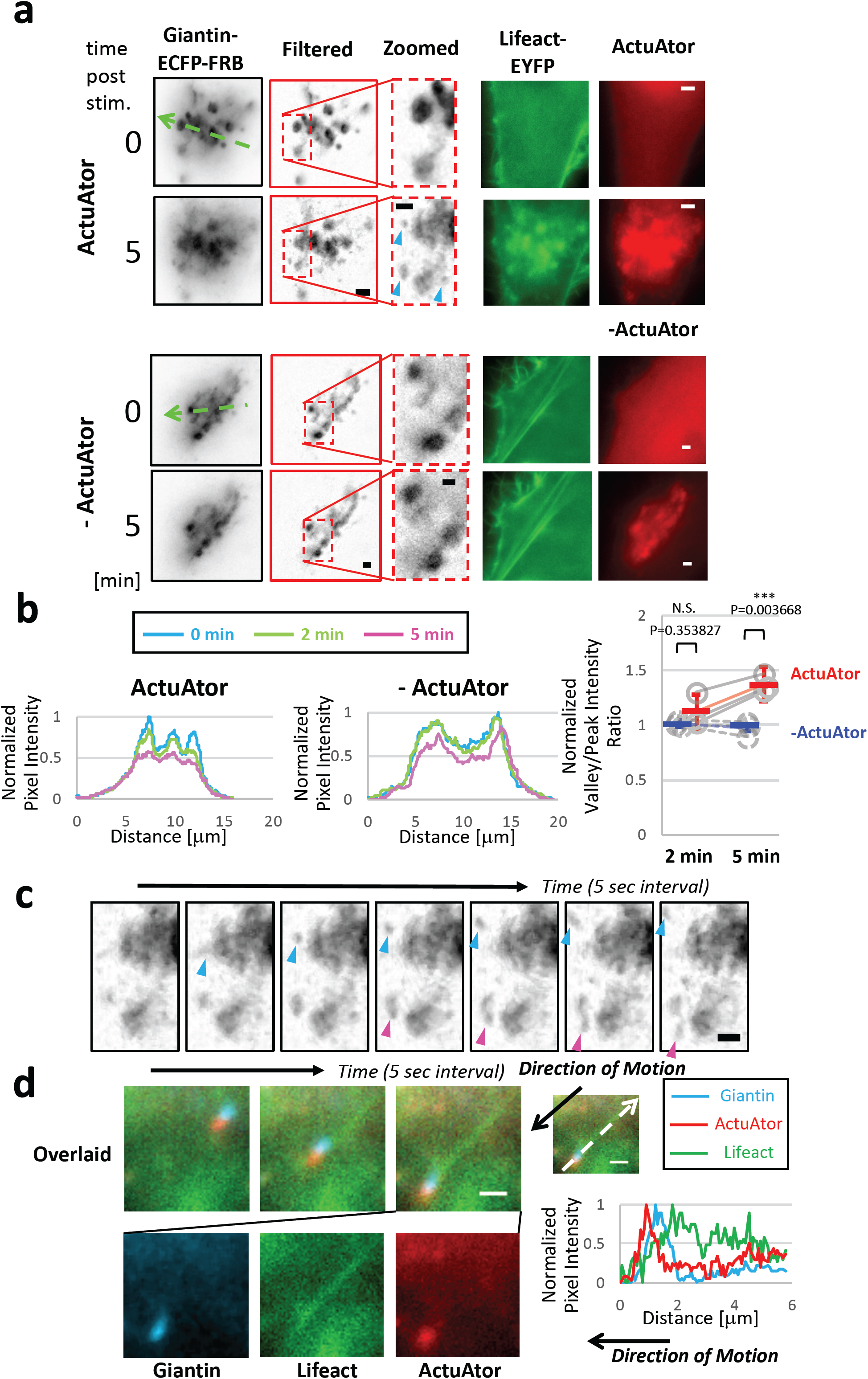
ActuAtor deforms Golgi apparatus, generating small moving fragments. a) Induced accumulation of ActuAtor peptide at Golgi apparatus induces change in Golgi morphology. In upper panels (ActuAtor), Golgi marker (giantin-ECFP-FRB, shown as inverted images), Lifeact (Lifeact-YFP, green), and ActuAtor (ActuAtor-FKBP-mCherry, red) images are shown right before (0 min) and five minutes after stimulus (5 min). Golgi morphology was extracted by filtering the giantin-ECFP-FRB images by top-hat morphology filter (Filtered, Zoomed). In lower panels (-ActuAtor), results without ActuAtor peptide is shown in similar fashion. Blue arrows in broken lines indicate the positions of linescan profiles shown in b). Scale bars: 1 μm in Zoomed images, 2 μm in other images. b) Left panels: Normalized fluorescence intensity profiles along the green arrows shown in a) are plotted at distinct time points (0, 2, 5 minutes after stimulus). Golgi morphology was significantly altered by ActuAtor recruitment, with intensity peaks abolished and blurred over time, while negative control peptide did not affect the profile. Right panel: Ratio between the intensity values at the peak and that at the valley in the linescan profiles are plotted. Results for each cell with (ActuAtor, N=3 cells) or without ActuAtor (-ActuAtor, N=3 cells) are plotted as grey circles with solid or broken lines, respectively. Mean ± standard deviation values are shown in red and blue markers for cells with or without ActuAtor, respectively. Statistical analysis was carried out with single-tailed Welch’s *t*-test. ***: p<0.01, N.S.: not significant. c) Sequential top-hat-filtered inverted fluorescence images of Golgi apparatus marker during ActuAtor-induced deformation. The data is the same as the one presented in upper panels in a). Smaller fragments highlighted by cyan and magenda arrowheads budded from larger entities and moved away. Interval between images are 5 sec. Scale bar: 1 μm. d) A Gogi apparatus marker-positive fragment traveling across the cytosol. Left panels; Cyan images are the Golgi marker, Giantin, green images are Lifeact, and red images are ActuAtor fluorescence images. Three sequential frames are shown as overlaid images. Interval between images are 5 sec. Note that the apparent displacement between Golgi marker (cyan) and ActuAtor (red) is due to the fast movement of the fragment during switching periods between distinct filter sets. Right panels; Normalized fluorescent intensity profiles along the broken-line white arrow in the overlaid image (upper left) for the three colors are shown as an overlaid plot (bottom). The direction of motion is shown as a black arrow in the upper left image, which is identical to the third frame in the left panels. Trailing comet of polymerized actin (Lifeact-positive) is clearly observed behind the fragment.

### ActuAtor induces deformation of nucleus

After its characterization at mitochondria and Golgi, ActuAtor was next targeted to other intracellular organelles, namely endoplasmic reticulum (ER) and outer nuclear envelope (ONE), using their targeting peptide, Sec61B, fused to FRB. Upon chemical dimerization, ActuAtor successfully accumulated at these subcellular locations where ECFP-FRB-Sec61B was localized, subsequently leading to three different types of deformation: patches, spikes, and finger-like protrusions (**Figure 4 a-c**). Actin-rich patches had a smooth boundary with round shapes often found in the central part of the nuclei (**Figure 4a, Supplementary movie S4**). Spikes had needle-like pointed morphology, which invaginates into nucleus typically initiated from the sides of the nuclei (**Figure 4b, Supplementary movie S5**). Finger-like protrusions were also invaginations from the sides, but with larger and round, less tapered shapes (**Figure 4c, Supplementary movie S6**). The three seemingly distinct phenotypes may actually be based on the same morphology but captured at a different angle on a microscope. Lastly, these morphological changes were never observed when FKBP alone (-ActuAtor) was used (**Figure 4d,e**).

**Figure 4.**
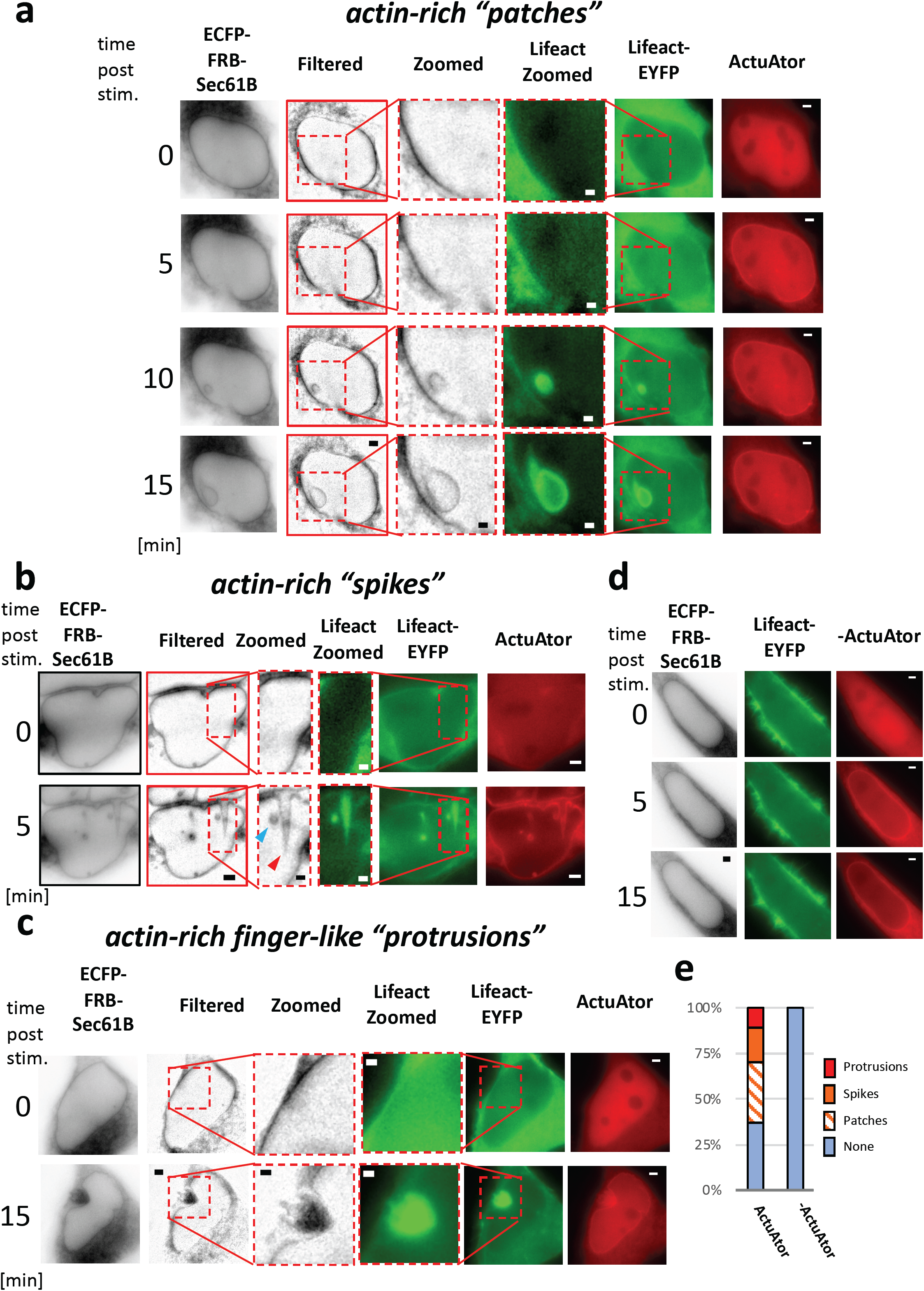
ActuAtor deforms outer nuclear envelope. ActuAtor recruitment to both endoplasmic reticulum and outer nuclear envelope leads to deformation of nucleus in some cells with varied phenotypes. These phenotypes were categorized into three classes according to the types of actin-rich structures observed: patches, spikes, and finger-like protrusions. a) A representative actin-rich patch found in deformed nucleus. Fluorescence images of ER and ONE marker (ECFP-FRB-Sec61B, shown as inverted images), Lifeact (Lifeact-EYFP, green), and ActuAtor (ActuAtor-FKBP-mCherry, red) at distinct time points after stimulus are presented. ONE morphology extracted by filtering the marker images by top-hat morphology filter is also shown as Filtered. Scale bars: 1 μm for Zoomed images, 2 μm in other images. b) Representative actin-rich spikes found in deformed nucleus. Needle-like pointed morphology characterizes the class of phenotype. Scale bars: 1 μm for Zoomed images, 2 μm in other images. c) A representative actin-rich finger-like protrusion found in deformed nucleus. Unlike patches, the invagination took place apparently from the side of the nucleus. Scale bars: 1 μm for Zoomed images, 2 μm in other images. d) Accumulation of fluorescently-tagged FKBP (-ActuAtor) at the nuclear envelope did not deform nucleus. Scale bars: 2 μm. e) Fraction of the cells categorized in each phenotypic class. N=27 cells and 22 cells for cells with (ActuAtor) or without ActuAtor (-ActuAtor), respectively.

### Form-function relationship of mitochondria probed by ActuAtor

Mitochondria undergo cycles of fission and fusion. The rate and frequency at which this dynamic cycle takes place primarily dictate overall morphology of mitochondria at a given time (Youle and van der Bliek, 2012; Friedman and Nunnari, 2014). Interestingly, the shape of mitochondria appears to correlate with their function (Picard et al., 2013; Pernas and Scorrano, 2016). Long, filamentous mitochondria exhibit upregulation of oxidative phosphorylation, while fragmented or vesicular mitochondria associate with compromised functionality. However, these observations are subject to complications. Genetic manipulation generally experiences a lag time of at least several hours between induction and final effect on the protein level. Several mitochondria proteins including Drp1 and Mfn1 have roles other than mitochondrial fission and fusion. A pharmacological inhibitor such as CCCP induces fragmentation of mitochondria, but it does so by interfering with a key mitochondrial function (i.e., maintenance of membrane potential), thus confounding its functional consequences in the light of its morphological change. ActuAtor-mediated acute deformation of mitochondria may offer an alternative, unique tool to probe the form-function relationship. Toward this end, we began by further characterizing the ActuAtor-induced mitochondria deformation.

Some vesicular-shaped mitochondria after deformation induced by ActuAtor had thin threads that appeared to bridge two mitochondria fragments (**Figure 1b,e, Figure 2**). We therefore evaluated the connectivity by asking if and how material transfer is achieved between the two deformed mitochondria by measuring the kinetics of fluorescence recovery after photobleaching (FRAP). As a marker, we used Su9-EYFP, a diffusive marker located within mitochondrial matrix. After mitochondrial deformation was induced by ActuAtor, FRAP measurement was conducted at a single spot within one of the mitochondria fragments under confocal microscope (**Figure 5a**). If the thin thread described above maintained connectivity, the Su9-EYFP fluorescence would eventually recover as the other mitochondria fragment will supply the proteins (**Supplementary Figure 4a**). When ActuAtor expressing cells were treated with DMSO as a control, Su9-EYFP fluorescence quickly recovered at the bleached spot (**Figure 5a,b**, top panels). CCCP is a small molecule drug that neutralizes mitochondria membrane potential which results in alteration of mitochondria morphology. These mitochondria appear to be severed into smaller pieces with no visible connections among them. When non-transfected cells were treated with CCCP and then underwent FRAP, we observed little to no fluorescence recovery as expected (**Figure 5a,b**, bottom panels). We then added rapamycin to ActuAtor-expressing cells, and observed very little fluorescence recovery, that is statistically indistinguishable from that of CCCP-treated cells (**Figure 5a,b**, middle panels, **Figure 5c, Supplementary Figure 4b**). This implies that mitochondria deformation by ActuAtor resulted in nearly complete isolation from each other, at least in terms of substance exchange between the mitochondria matrices.

**Figure 5.**
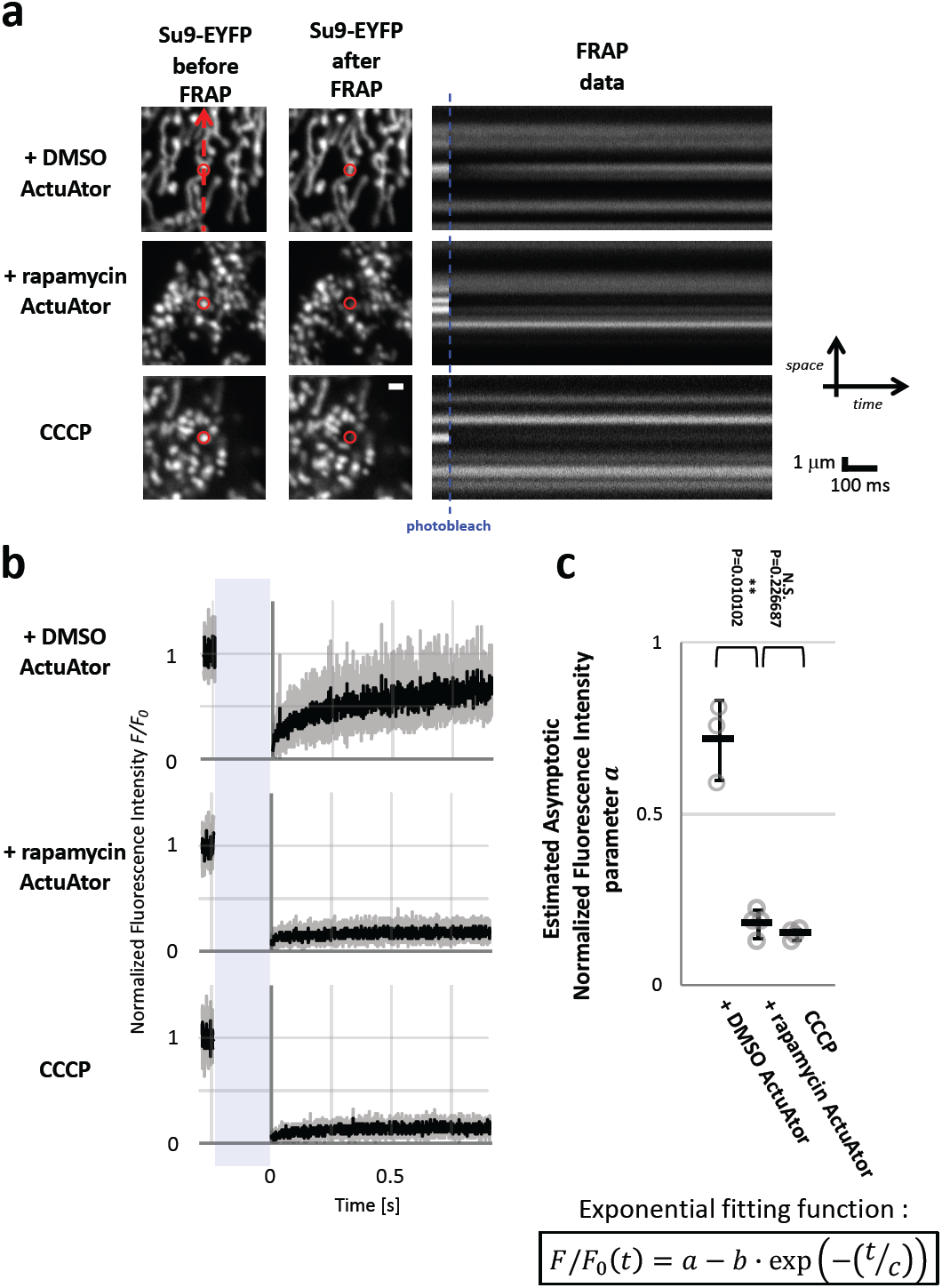
Characterization of ActuAtor-deformed mitochondria by FRAP measurement reveals abolishment of connectivity. a) Left columns: Two-dimensional *xy*-scanned images of Su9-EYFP fluorescence before and after FRAP measurement in each condition are shown. Photobleaching was done at a single point depicted by red circles in the images. Laser scanning during FRAP measurement was carried out along vertical lines with the photobleaching spot in the center, as the red broken arrow in the left upper image. Fluorescence intensity is significantly reduced after FRAP in experiments with ActuAtor translocation (+rapamycin ActuAtor, N=4 cells) or CCCP treated cells (CCCP, N=4 cells), while it recovered in cells without ActuAtor translocation (+DMSO ActuAtor, N=3 cells), which is consistent with apparently low connectivity between deformed mitochondria in former cases. Scale bar: 1 μm. Right column: Results of FRAP measurement shown as *xt* space-time presentation. b) Normalized fluorescence intensity at photobleached spot plotted against time for each condition. Black line is the mean value, while gray transients are mean ± standard deviation. Period of photobleaching is shown as pale blue shade in each plot. c) Asymptotic recovery of normalized fluorescence intensity quantified by regression analysis is plotted for each condition. FRAP data was fitted to the formula shown in the lower panel. Fitted value of asymptotic recovery value *a* is shown in the plot. Fitted values of other parameters are shown in Supplementary Figure 5. Single-sided Welch’s *t*-test was used for analysis. Connectivity between ActuAtor-deformed mitochondria estimated by the current FRAP analysis was thus indistinguishable from that between mitochondria fragmented by CCCP treatment. **: p<0.05, N.S.: not significant.

After validating the basic properties of ActuAtor-deformed mitochondria, we next examined if and how the mitochondrial morphology and function are interlinked by quantitatively monitoring key indicators of mitochodnria functions as they underwent deformation. These include ATP concentration (**Figure 6a,d**), membrane potential (**Figure 6b,e**), and concentration of reactive oxygen species (ROS) (**Figure 6c,f**), using mitochondria-targeted ATeam 1.03 (mitAT1.03) (Imamura et al., 2009), tetramethylrhodamine ethyl ester (TMRE) (Ward, 2010; Chazotte, 2011), and mitochondria-targeted HyperRed (HyperRed-mito) (Ermakova et al., 2014), respectively. Time-lapse fluorescence measurement was carried out for 15 minutes (5 minutes pre-stimulus and 10 minutes post-stimulus) in cells cultured in either glucose- or galactose-containing culture medium. Replacement of glucose with galactose makes cells shift their energy source from cytosolic glycolysis to mitochondrial oxidative phosphorylation. To evaluate cell-type dependency, two cell types, HeLa and COS-7, were adopted. In all conditions tested, positive controls (**Figure 6a,d**, orange lines and plots) were designed according to previous studies, and along with negative controls, were used to evaluate the sensitivity of each experiment.

**Figure 6.**
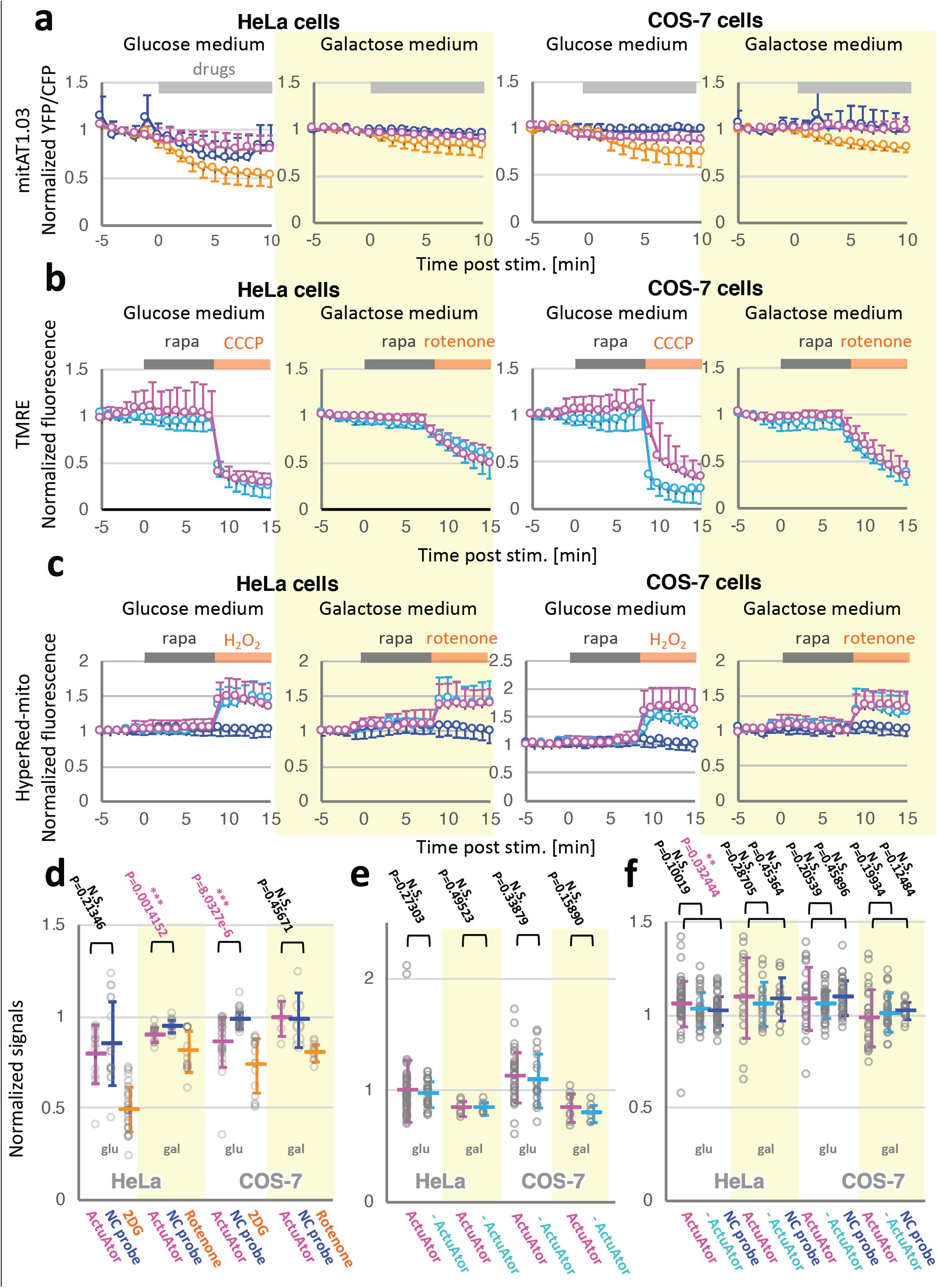
Mitochondria deformation by ActuAtor shows modest effects on mitochondrial functions. Effects of mitochondria deformation on mitochondria functions were evaluated by measuring three distinct parameters: ATP concentration at mitochondria, membrane potential across inner mitochondrial membrane, and reactive oxygen species (ROS) concentration at mitochondria. Each property was examined by fluorescence imaging using suitable indicator, mito-ATeam1.03 for ATP, TMRE for membrane potential, and HyperRed-mito for ROS. Two cell types, HeLa and COS-7 were used for the measurement. a) ATP concentration measurement by mitAT1.03 at mitochondria in conjunction with mitochondrial deformation in HeLa cells and in COS-7 cells. Results with mitochondria deformation by *ActuAtor* (magenta), the same condition with ATP-insensitive mutant probe (blue), and positive control where mitochondrial function was abolished by drug treatment (orange) are shown as mean ±standard deviation. The right panels highlighted by yellow background are the results in cells cultured in galactose medium. As positive controls, CCCP and rotenone were adopted for experiments in cells cultured in normal or glucose medium and those cultured in galactose medium, respectively. N=12,17,31 cells for HeLa cells with ActuAtor, ATP-insensitive probe, and positive control, N=26,34,16 cells for COS-7 cells, respectively, in glucose media conditions. N=15,13,16 cells for HeLa cells and N=6,6,11 cells for COS-7 cells, respectively, in galactose conditions. b) Membrane potential measurement by TMRE. TMRE fluorescence was monitored at mitochondria before and after ActuAtor-induced deformation, followed by adding positive control drugs to the medium. Results with galactose medium are highlighted by yellow background. N=43 and 31 cells with or without ActuAtor in HeLa cells, N=35 and 19 cells in COS-7 cells, respectively, in glucose conditions. N=7 and 6 cells for HeLa cells, N=9 and 7 cells for COS-7 cells, respectively, in galactose conditions. c) ROS concentration measurement by HyperRed-mito. After deformation by ActuAtor, hydrogen peroxide (H2O2) or rotenone was added to the cells cultured in glucose or galactose media, respectively, as positive control conditions. Results in galactose medium-cultured cells are highlighted by yellow background. N=53,29,53 cells with or without ActuAtor and with negative control probe in HeLa cells, N=23,43,34 cells for COS-7 cells, respectively, in glucose conditions. N=18,17,14 cells for HeLa cells, N=26,26,15 cells for COS-7 cells in galactose conditions. d-f) Statistical analyses on normalized signals ten minutes after stimulus. Results for mitochondrial ATP concentration (d), membrane potential (e), and mitochondrial ROS concentration are shown. Data points from individual cells are plotted as grey circles, while mean ± standard deviation values are shown in markers. In all plots, magenta markers are the results with ActuAtor. Results of cells cultured in galactose medium are highlighted by yellow backgrounds.

For ATP measurement, we observed a modest decrease in ActuAtor-deformed mitochondria under two out of four conditions, HeLa cells in galactose medium and COS-7 cells in glucose medium (**Figure 6d**). The remaining conditions did not show any change in ATP concentration at deformed mitochondria. One caveat in interpreting these data would be that actin polymerization induced by ActuAtor is expected to consume ATP in the vicinity of mitochondria, which may interfere with measurement of ATP produced by mitochondria. Additional metabolic measurement may complement this potential pitfall.

For membrane potential measurement, we did not observe a significant decrease at ActuAtor-deformed mitochondria under any condition tested (**Figure 6b,e**). There was no clear tendency of the signal being reduced, judging from the mean values. The type of culture media did not affect the results.

With ROS concentration measurement, we performed experiments under eight conditions, along with statistical tests using two negative controls (**Figure 6c,f**). As a result, there was only one condition, HeLa cells in glucose medium, that exhibited a statistically significant change. However, the extent of this change was modest (less than 10%).

To summarize, by monitoring mitochondria functions in real time before and after deformation, we found almost no change under the majority of conditions we examined. There were three conditions out of 12 where changes were observed, albeit modest. We could not find a clear pattern as to the relationship between their morphology and functions, at least under these experimental conditions.

### Deformed mitochondria yielded to mitophagy-dependent degradation

Although we did not detect striking functional effects of *ActuAtor*-dependent mitochondrial deformation, there is a possibility that altered morphology affects other mitochondrial properties such as turnover of the organelle over a longer time period. Mitophagy, which is autophagy-dependent degradation of mitochondria (Pickles et al., 2018), is a promising candidate, because hyperfused mitochondrial morphology due to Drp1 knockout has been linked to reduced mitophagy, which was rescued by reversing the morphology via additional knockout of a fusion protein, Opa1 (Yamada et al., 2018). It is therefore plausible that the more fragmented morphology observed with ActuAtor would increase the flux of mitochondria degradation toward mitophagy.

To test this, we employed Parkin as a mitophagy marker. Parkin is recruited to the surface of mitochondria destined to undergo mitophagy. HeLa cells supplemented with exogenous expression of Parkin have been used as a valid experimental model for Parkin-dependent mitophagy. Using this system, we evaluated colocalization between EYFP-Parkin and ActuAtor-deformed mitochondria as a readout of Parkin-dependent mitophagy. Twenty-four hours after the ActuAtor operation, we observed a modest increase in the colocalization value (2-3%), while the negative control with FKBP alone (-ActuAtor) showed no increase, and the positive control of CCCP treatment induced 10% increase (**Supplementary Figure 5a,b**). Our result suggests that fragmented, smaller mitochondria are more susceptible to mitophagy. One caveat is that the observed long-term effect may not be simply attributed to the difference in size or shape of mitochondria, as unexpected lateral effects of induced actin polymerization may become significant over time.

### ActuAtor disperses non-membrane-bound structures

ActuAtor was demonstrated to deform intracellular structures with surrounding membranes. We further tested ActuAtor for another class of intracellular structures known as non-membrane-bound organelles or biomolecular condensates (Banani et al., 2017; Wheeler and Hyman, 2018). They are structures composed of biomolecules, most typically proteins and RNAs, including RNA granules proposed to be involved in cellular processes such as DNA transcription and RNA translation, as well as in pathogenesis of neurodegenerative diseases(Weber and Brangwynne, 2012). While there are many techniques available to induce assembly of biomolecules to form artificial granules(Nakamura et al., 2018), physiological roles of native granules remain largely unknown, due to a lack of techniques to manipulate the granules in their intact form.

To assess whether ActuAtor could interfere with organization of native RNA granules, we chose stress granules as a benchmark. Stress granules are mRNA-containing granules that are transiently formed in cells under various stresses (Kedersha et al., 2013; Protter and Parker, 2016). They are also associated with neurodegenerative diseases such as amyotrophic lateral sclerosis (Zhang et al., 2018). However, the physiological functions of stress granules are still under active debate. We thus performed the ActuAtor operation in cells with stress granules that were formed in response to prior treatment with sodium arsenite that serves as a chemical stress. We first made an FRB fusion protein with a stress granule marker TIA-1, which was co-expressed with chemically-inducible ActuAtor. Upon addition of rapamycin, ActuAtor was efficiently recruited to the stress granules, followed by actin polymerization (**Figure 7a, Supplementary movie S7**). Actin polymerization appeared to occur throughout the individual stress granules, which may simply reflect the distribution of TIA-1. As a result, stress granules were dispersed over time, despite the presence of sodium arsenite throughout the imaging duration. However, when a negative control peptide FKBP alone with no ActA portion (-ActuAtor) was accumulated at the stress granules, we unexpectedly observed moderate dispersion of the granules (**Supplementary Figure 6a**). Although the frequency of dispersion was significantly lower compared to ActuAtor, this background effect could be problematic in assessing physiological functions of stress granules. We therefore replaced TIA-1 with two other stress granule markers, G3BP1 and FXR1b, as an anchor for the ActuAtor recruitment, and characterized their efficiency of induced dispersion. When either FRB fusion protein, CFP-FRB-G3BP1 or CFP-FRB-FXR1b, was co-expressed in cells with ActuAtor, rapamycin addition led to accumulation of ActuAtor at the stress granules, followed by their striking dispersion (**Figure 7b**). Quantification of the dispersion obtained with each of the three anchor proteins highlighted difference in dispersion kinetics (red lines, **Figure 7c**) with a descending order of TIA-1, G3BP1 and FXR1b. We also analyzed dispersion kinetics using a negative control probe, FKBP alone (i.e., -ActuAtor), which also indicated variations (blue lines, **Figure 7c**). Compared to TIA-1, G3BP1 induced slower dispersion, while FXR1b triggered almost no dispersion. Since G3BP1 and FXR1b exhibited the less significant background effect compared to TIA-1, we further characterized these by monitoring the morphology of stress granules as they underwent ActuAtor-mediated dispersion. We found a qualitative difference; while G3BP1 uniformly dispersed across a stress granule with an increasingly smoother edge, FXR1b broke into smaller fragments that were clearly visible under the microscope (**Figure 7d**). The distinct kinetics and dynamics of ActuAtor-induced stress granule dispersion could be attributed to the differential distribution of these anchors within granules and/or to the distinct effects on biophysical properties of granules due to the protein overexpression. Nevertheless, we demonstrated that ActuAtor can disperse native stress granules in living cells.

**Figure 7.**
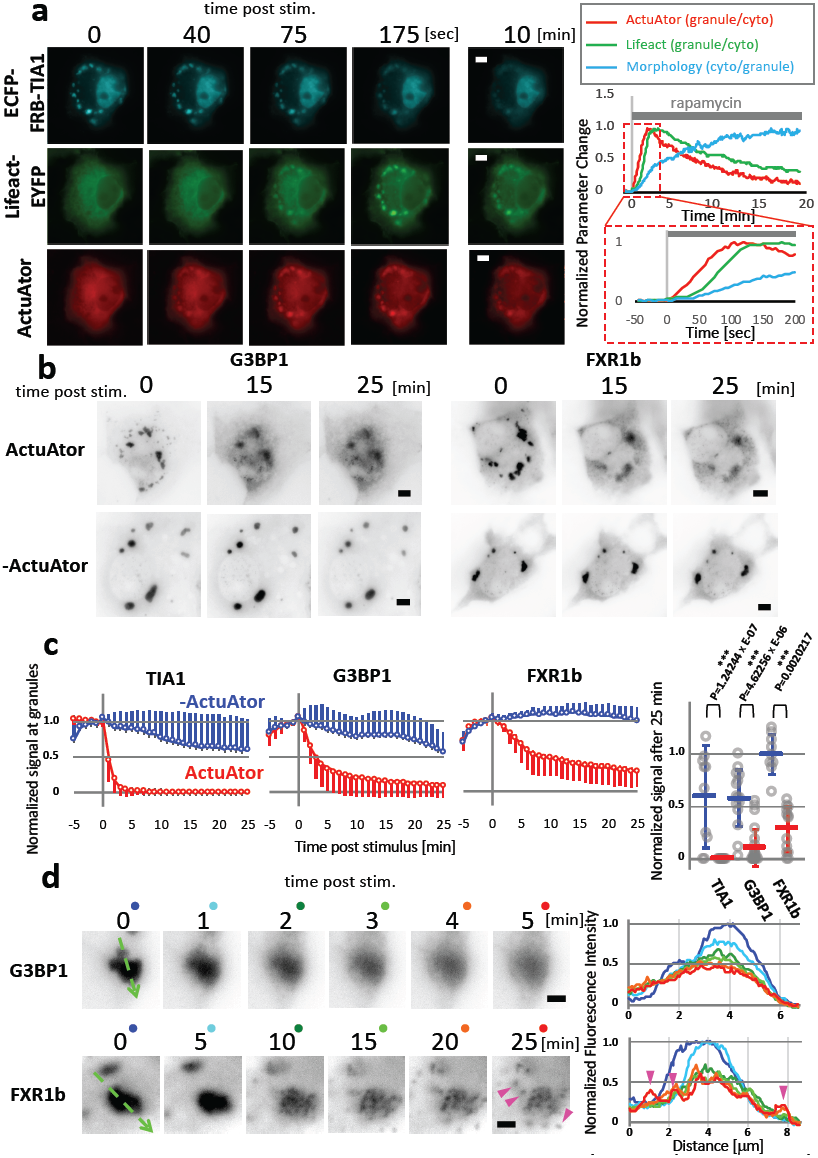
ActuAtor disperses a non-membrane-bound organelle, stress granules, in an inducible manner. a) Stress granules were dispersed by Actuator recruitment, preceded by actin polymerization at the site. Fluorescence images of stress granule morphology marker (ECFP-FRB-TIA1, cyan), Lifeact (Lifeact-EYFP, green), and ActuAtor peptide (ActuAtor-FKBP-mCherry) at distinct time points are shown in the left panel. In the right panel, Normalized ratio of fluorescence intensities of ActuAtor (red), Lifeact (green) and ECFP-FRB-TIA1 (cyan) at the granules to those in the cytosol are plotted against time. Enlarged plot is shown below in red broken-line rectangle for earlier time points. b) Multiple stress granule markers can be used to recruit ActuAtor for inducible dispersion of granules. Stress granule morphology as inverted fluorescence intensity images of an independent stress granule marker, EYFP-PABP1, are shown. Two stress granule markers, G3BP1 and FXR1b, were used to recruit ActuAtor to the granules, as ECFP-FRB-G3BP1 and ECFP-FRB-FXR1b, respectively. Both markers recruited ActuAtor successfully, resulting in significant stress granule dispersion (ActuAtor), compared to negative control (-ActuAtor). Scale bars: 5 μm. c) Left panel: Temporal evolution of integrated fluorescence signal of a marker, EYFP-PABP1 at the granules with (ActuAtor, red) or without ActuAtor recruitment (-ActuAtor, blue). Normalized signals are plotted for three stress granule markers, TIA-1, G3BP1, and FXR1b, used to inducibly accumulate the peptides. N=10 regions from 5 cells (-ActuAtor, TIA1), 10 regions from 5 cells (ActuAtor, TIA1), 15 regions from 12 cells (-ActuAtor, G3BP1), 23 regions from 12 cells (ActuAtor, G3BP1), 9 regions from 5 cells (-ActuAtor, FXR1b), and 13 regions from 6 cells (ActuAtor, FXR1b). Right panel: Normalized integrated signals 25 minutes after stimulus are plotted for distinct conditions. Results in individual cells are plotted as grey circles, while mean ± standard deviation values are shown in red and blue markers for cells with or without ActuAtor. d) Difference in stress granule morphology during the dispersion by ActuAtor recruited by distinct marker proteins. Inverted EYFP-PABP1 fluorescence images in a representative region at distinct time points are shown for each condition, where ECFP-FRB-G3BP1 (G3BP1) or ECFP-FRB-FXR1b (FXR1b) was used to recruit ActuAtor. In the right panel, line-scan profile of the fluorescence intensity along green broken-line arrows are presented in colors corresponding to those of dots shown besides images in the left panel. FXR1b-dependent recruitment resulted in more distinguishable fragments, highlighted by magenta arrowheads in both panels. Scale bars: 2 μm.

## Discussion

We report the development and implementation of ActuAtor, a molecular tool that can generate force in living cells for acute deformation of intracellular structures such as mitochondria, Golgi, nucleus and stress granules (**Supplementary Figure 6b**, for a summary illustration). ActuAtor possesses several unique characteristics: genetically encoded nature, modular design, and remote control (using light or chemical). ActuAtor can be introduced into cells and tissues by simple transfection or viral infection of its encoding DNA, or by conventional genetic manipulation. Due to the modular design, ActuAtor can be targeted to arbitrary subcellular structures, as long as their targeting peptides are available. ActuAtor can be operated without directly touching the cells; it offers a means to assessing mechanobiological events in intact cells. ActuAtor can also be controlled by at least two stimuli, namely a chemical dimerizer and light. Both CID and LID have orthogonal pairs of dimerizing peptides, with which we could potentially multiplex ActuAtor operation for more demanding manipulation. Since virtually all cells expressing ActuAtor undergo its operation, ActuAtor should be compatible with a large scale experiment such as a metabolomics analysis.

As an emerging technique, there is room for improvement. The magnitude of force exerted by ActuAtor can be modified by varying the concentration of rapamycin and/or expression level of ActuAtor and FRB anchors, as well as their ratio. Regardless, it is currently challenging for ActuAtor to exert a pre-determined amount of force against a target object. Another potential drawback is the reliance on actin polymerization, a fundamental and prevailing biological process. While no noticeable effect on endogenous actin stress fibers was detected (**Supplementary Figure 2e**), ActuAtor operation could interfere with other actin-related structures. It would be ideal if we could design a molecular tool for force generation based on a building block that is orthogonal to biological processes. Nevertheless, it is imperative to design appropriate control experiments for the use of ActuAtor.

In probing the form-function relationship of mitochondria, we observed that ActuAtor-deformed mitochondria could still perform representative mitochondria functions almost normally. Based on this, we conclude that mitochondria morphology has only marginal effects on these functions. In other words, it appears that mitochondria morphology follows its functions, but not vice versa. This notion challenges the commonly-accepted form-function relationship of mitochondria. However, we cannot exclude the possibility that other aspects of mitochondrial biology may have been affected in response to the morphological changes. Functions remaining to be tested include oxygen consumption, apoptosis induction, lipid metabolism, Parkin-independent mitophagy, and trafficking along microtubules. It is also exciting to explore form-function interplay of other organelles, for example, by addressing how the nucleus senses external mechanical stress to control gene expression for cell fate determination, and how plasma membrane tension regulates cell migration properties.

Another exciting utility of ActuAtor is dispersion of stress granules. Considering the common physical mechanism shared across diverse non-membrane-bound organelles, and due to the modular nature of ActuAtor design as mentioned earlier, it is probable that ActuAtor could be adapted to disperse other biomolecular condensates such as Cajal bodies, P granules, P-bodies, and early-stage inclusion bodies found in neurodegenerative diseases. Readily generalizable loss-of-function paradigm using ActuAtor can thus be a promising tool to start revealing their physiological and pathophysiological functions. Importantly, the precise mechanism of the dispersion has yet to be understood. Based on the slightly distinct observations with the three different stress granule markers, we speculate that both force generation and dense actin network formation within the granules contribute to weakening association between protein and RNA constituents. Nevertheless, more thorough characterization is needed to fully appreciate the molecular mechanism underlying stress granule dispersion.

In summary, ActuAtor is the first tool that can exert force to deform an arbitrary intracellular structure in living cells. The tool is genetically encoded and is actuated by chemicals or light, which makes ActuAtor compatible with *in vivo* applications, as has been demonstrated by optogenetic tools such as channelrhodopsins. Therefore, ActuAtor enables future elucidation of the currently underexplored field, “intracellular mechanobiology”.

## Acknowledgements

The authors thank the following researchers for sharing research reagents: Hiroomi Imamura for ATeam1.03 constructs; Vsevolod Belousov for HyperRed constructs; Nancy Kedersha for stress granule-related constructs; Charlotte Sumner for 50B11 cells; and Hiromi Sesaki for Drp1 knockout cells and Parkin construct. Our appreciation is extended to Erin Goley, Jodi Nunnari and Nancy Kedersha for insightful discussions and comments, Brett Andrew McCray for technical advices on experiments, and to our lab members including Brian Huang and Robert DeRose for experimental assistance and fruitful discussions. We acknowledge support from the National Institutes for Health (5R01GM123130 to T.I.), and the DoD DARPA (HR0011-16-C-0139 to T.I.).

## Author Contributions

H.N. conceived the research. H.N. and T.I. designed experiments and wrote the manuscript. H.N. performed the majority of experiments and data analysis thereof. E.R. performed some of the mitochondria experiments, and D.D. performed some of the stress granule experiments, both under the supervision of H.N. and T.I. H.M. performed and analyzed *in vitro* experiments using proteins purified by S.R..

## Methods

### Cell Culture and Chemical Reagents

HeLa cells, COS-7 cells, NIH3T3 cells, and U2OS cells were obtained from ATCC, and cultured in Dulbecco’s modified Eagle’s Medium (corning, 10013CV) supplemented with 10 % fetal bovine serum (FBS) (Sigma Aldrich, F6178) at 37 °C, 5% CO_2_. 50B11 cells were kindly provided by Charlotte Sumner, and cultured in Neurobasal medium (Thermo Fisher, 21103049) supplemented with 5% fetal bovine serum, 0.2 % v/v B27 (Thermo Fisher, 17504044), 0.1% v/v Glutamax (Thermo Fisher, 35050061), penicillin-streptomycin (Thermo Fisher, 15140163), and 0.2% w/v glucose. 50B11 cells were treated with 50 μM forskolin (Millipore Sigma, F6886) in 0.2% fetal bovine serum-containing Neurobasal medium 24 hours prior to imaging, allowing the cells to change morphology into elongated, neuron-like shape with long processes. Drp1 knockout mouse embryonic fibroblasts (MEFs) were kind gift from Hiromi Sesaki. Drp1 MEFs were cultured in 10% FBS-containing Iscove’s modified Dulbecco’s medium (Gibco, 12440053).

### DNA Plasmids

All the plasmids were constructed by standard subcloing techniques based either on pECFP, pEYFP, and pmCherry plasmids. ActA sequences were synthesized with codon usage optimized for mammals (Genscript). Signal sequences for specific targeting to organelles were synthesized as oligo DNAs and subcloned. TIA-1, PABP-1, G3BP1, and FXR1b sequences are PCR-amplified from constructs kindly provided by Nancy Kedersha. EspFu5R sequence and Nck SH3 sequence are provided by Wendell Lim and Matthew Kennedy, respectively, and PCR amplified for subcloning. ATeam1.03 plasmids and HyperRed constructs are kindly provided by Hiroomi Imamura and Vsevolod Belousov, respectively. Major plasmids used in the manuscript will be deposited and available from Addgene.

### 6xHis-ActA (1-584)-FRB-CFP purification

ActA(1-584)-FRB-ECFP was subcloned in pET28a plasmid to have N-terminally tagged with 6x His tag for purification. Inoculated a colony of BL21-CodonPlus(DE3)-RIL competent cells (Agilent Technologies 230245) transformed with the plasmid DNA in 10 ml of LB media supplemented with kanamycin and grew overnight at 37°C, 220 rpm. Transferred the culture to 1L of LB/kanamycin and expanded at 37°C, 220 rpm till OD_600_ reached ∼0.5. Induced the cells with IPTG at a final concentration of 0.4 mM and continued culture at 19°C, 220 rpm, overnight. Pelleted the cells at 4500 × g, 4°C for 15 min. Resolubilized the pellet in the lysis buffer (10 mM Tris pH 7.6, 100 mM NaCl, 10 mM imidazole, 10% glycerol, 1 mM β-mercaptoethanol and cOmplete™ EDTA-free protease inhibitor tablet (Sigma Aldrich 11836170001)). Lysed the cells with a microfluidizer. Harvested the soluble fraction by centrifugation at 16,000 x g, 4°C for 40 min. Filtered the supernatant with 0.2 μm syringe filters (Thermo Fisher F25006). Equilibrated a 5 ml His-Trap HP column (GE Life Sciences) with the lysis buffer lacking protease inhibitors. Eluted the protein using a linear imidazole gradient from 50-500 mM over 20 column volumes. The eluted fractions were analyzed on a SDS-PAGE gel and subsequently the peak fractions detected on the Coomassie blue (Biorad 1610400) stained gel were pooled and dialyzed in a 10 kDa SnakeSkin™ dialysis tubing (Thermo Fisher 68100) in 2L of storage buffer (10 mM Tris-HCl pH 7.6, and 50 mM NaCl, 10% glycerol, and 1 mM β-mercaptoethanol) at 4 °C, overnight. The next day the protein was dialyzed for an additional 3 hours in fresh storage buffer and subsequently applied to a Superdex 200 10/300 GL (GE Life Sciences) equilibrated in the same buffer. The peak fractions were combined, and concentrated with an Amicon® Ultra 10 kDa centrifugal filter unit (Millipore UFC801024). Aliquoted protein vials were flash-frozen and stored at −80 °C.

### In Vitro Pyrene Assay of Actin Nucleation Efficiency

Rabbit skeletal muscle actin (Cytoskeleton, Inc. AKL99), pyrene-labeled actin (Cytoskeleton, Inc. AP05), Arp2/3 complex (Cytoskeleton, Inc. RP01P), and GST-WASP-VCA (Cytoskeleton, Inc. VCG03-A) were reconstituted and used as described in manufacture’s protocols except for size exclusion chromatography (GE Superdex 200 increase 10/300) purification of actin. Pufied actin was stored at 4°C with dialysis against G-buffer 1 (2 mM Tris-HCl (pH 7.5 at rt), 0.1 mM CaCl_2_, 0.2 mM ATP, 0.5 mM DTT, 1 mM NaN_3_).

Pyrene-actin assay was performed by using FluoroMax 3 and polymerization kinetics were measured by using FluoroMax 3 operated by Datamax software (HORIBA, 365 nm emission 1 nm bandwidth and 497 nm excitation 5 nm bandwidth). Proteins for pyrene assay were prepared as follows. Actin and pyrene-labeled actin were diluted with G-buffer 2 (5 mM Tris-HCl (pH 7.5), 0.2 mM CaCl_2_) and mixed to make 5% pyrene labeled 4 µM actin solution. Arp2/3, Actuator-FRB-ECFP, and GST-WASP-VCA were diluted F-buffer (10 mM Tris-HCl (pH 7.5 rt), 50 mM KCl, 2 mM MgCl_2_, 1 mM ATP). Reaction was started by mixing 28 µL of G-buffer 2, 12.5 µL of 4 µM actin (5% pyrene-labeled), 4.5 µL 10xF buffer, and 5 µL of 100 nM Arp2/3. ActuAtor, GST-WASP-VCA, or F-buffer (negative control) was added 5–10 minutes after measuring baseline.

### Transfection of Cultured Cells and Live-cell Imaging of Organelle Deformation

Cells were seeded on a LabTek 8-well chamber (Thermo Scientific) coated with poly-D-lysine hydrobromide (Sigma Aldrich, P6407) solubilized in sterile water at 0.1 mg/ml. Transfection was done using X-tremegene9 transfection reagent (Roche, 6365779001) following manufacturer’s instruction, unless specified. Ratio amount between plasmids used for transfection was adjusted so that plasmids coding ActuAtor were relatively less (10-50% of organelle or stress granule markers), which resulted in efficient translocation of the ActuAtor peptides. Organelle markers or signal sequences used for translocation to mitochondria, Golgi apparatus, and outer nuclear envelope/endoplasmic reticulum were derived from Tom20, Giantin, and Sec61B proteins, respectively. As an exception, mitochondria translocation in 50B11 cells were carried out with signal sequences of MoA protein, as Tom20 signal sequence was poorly expressed in the cell type for unknown reason.

Cells were imaged 24 hr after transfection. For live-cell imaging of organelle deformation, Eclipse Ti inverted fluorescence microscope (Nikon) equipped with x60 oil-immersion objective lens (Nikon) and Zyla 4.2 plus CMOS camera (Andor) was used, unless specified. The system was controlled by NIS element software (Nikon). Stage-top incubator (Tokai Hit) was used to incubate samples at 37 °C, 5 % CO_2_ during imaging. For chemical stimulation, 100 μM rapamycin (LC Laboratories, R-5000) stock in DMSO was diluted in extracellular medium and used at final concentration of 100 nM. Interval between consecutive frames were either five or ten seconds.

### Quantification of Mitochondria Deformation

A region with readily observable mitochondria morphology was chosen as a region of interest (ROI) in each cell analyzed. Fluorescence images of mitochondria marker within ROIs were then processed by “top-hat morphology filter” in MetaMorph software (Molecular Devices). Morphology filtered images were binarized, and skeletonized using Fiji software (Image J). The skeletonized mitochondria morphology was analyzed by “analyze skeleton” menu of the software, and “the length of the longest shortest path” was extracted for each mitochondrion. The resultant lengths were averaged over each ROI to obtain mean mitochondria lengths in the formula shown in Figure 1c, using mitochondria images within the same region before and after stimuli.

### Confocal Live-cell Imaging and Optogenetic Stimulation

Light-inducible version of ActuAtor was imaged with LSM800 confocal microscope (Zeiss) using 514 nm and 561 nm lasers to excite EYFP and mCherry, respectively. To induce dimerization between iLID and SspB, 488 nm laser was used taking advantage of bleaching function in Zen Black software (Zeiss). In order to avoid background dimerization, 514 nm laser intensity was kept as low as possible during EYFP imaging. Samples were incubated at 37 °C, 5 % CO_2_ using built-in stage-top incubator during imaging.

### FRAP Measurement and Analysis

FRAP measurement of Su9-EYFP was done by line-scan mode of LSM800 confocal microscope and Zen Black software (Zeiss) to observe the fast recovery kinetics after photobleaching. Bleaching function of the same software was used for the photobleaching with 514 nm laser, which was also used for EYFP imaging at lower intensity. Photobleaching aimed at a single spot located in the middle of the line across which the laser scanned for imaging. To analyze the data, a Region of three pixels in width, which contained the photobleached spot in the middle, was used as a ROI. Each transient of integrated intensity over the ROI was fitted to the exponential function shown in Figure 5c, using R software (the R Foundation), to obtain three fitting parameter values, *a, b*, and *c*.

### Fluorescent Biosensor Imaging of Mitochondria Functions

Cells were seeded on a coverslip coated by poly-D-lysine hydrobromide and cultured in 6-well culture plate. All experiments were performed with two groups of cells cultured in distinct media for 48-72 hours prior to the imaging; glucose medium and galactose medium. We here refer to the conventional DMEM containing 25 mM glucose as glucose medium, while galactose medium contained 10 mM galactose in place of glucose. It has been reported that cells cultured in glucose medium metabolize majority of glucose via glycolysis, while in galactose medium, ATP synthesis through citrate cycle and mitochondrial respiration rate are increased (Reitzer et al., 1979; Rossignol et al., 2004). Imaging was performed with an inverted fluorescence microscope, Axiovert 135 TV (Zeiss) equipped with a QIClick charge-coupled device camera (QImaging).

For ATP measurement in mitochondria matrix, mitochondria-targeted FRET-based ATP biosensor, mitAT1.03 was used (Imamura et al., 2009). Transfection was performed in the 6-well culture plate using FuGene-HD (Promega, E2311) following manufacturer’s instructions. Fluorescence was measured by drawing three circular ROIs at mitochondria and 3 ROIs for background fluorescence. The net fluorescence was calculated by taking the difference of the averages. Methods for Analyses of FRET ratio change are described in previous reports (Miyamoto et al., 2015, 2018). In glucose medium condition, 25 mM of 2-deoxy-D-glucose was used to lower ATP level as a control. In galactose medium condition, 500 nM rotenone was used instead.

To measure membrane potential across mitochondria inner membrane, tetramethylrhodamine ethyl ester (TMRE) staining dependent on the membrane potential across the membrane was used. Staining was performed at 7 nM final concentration of TMRE for HeLa cells, while 35 nM TMRE was used for COS-7 cells. As a control, a proton ionophore, carbonyl cyanid m-chlorophenyl hydrazone (CCCP) was used to abolish the membrane potential at final concentration of 10 μM. Instead of ActuAtor containing mCherry used in majority of experiments described in the current report, we used EYFP-labeled ActuAtor in TMRE and HyperRed experiments described below.

In ROS concentration measurement experiments, we used a mitochondria matrix-targeted ROS biosensor, HyperRed-mito (Ermakova et al., 2014). As a control, 10 mM H_2_O_2_ was used for glucose medium experiments, while 500 nM rotenone was used for galactose experiments. As a negative control peptide, C199S mutant of HyperRed-mito (Ermakova et al., 2014), which we denoted as NC probe, was used.

### Stress Granule Dispersion

Live-cell imaging of stress granule dispersion was performed in COS-7 cells, with the same experimental setups for organelle deformation experiments described above, while longer interval between frames of one minute was adopted according to the slower kinetics observed. To form stress granules, cells were treated with 0.5 mM sodium arsenite contained in culure medium for 30 minutes prior to the imaging. During imaging, cells were incubated in the stage-top incubator in the sodium arsenite-containing culture medium. As a stimulus, rapamycin was manually added at the final concentration of 100 nM.

To quantify the dispersion induced by ActuAtor, fluorescence intensity of an independently co-expressed stress granule marker, EYFP-PABP1 was used. Regions with typical stress granules were selected as ROIs in the first frame for the analysis on MetaMorph software. Threshold was then manually adjusted to define the area of granules in the same frame. EYFP-PABP1 fluorescence images in all frames were binarized based on the threshold to create binarized masks. Cytosolic EYFP-PABP1 fluorescence intensity outside granules was also defined as a background level for subtraction. Background-subtracted fluorescence intensity was then integrated over stress granule area based on the mask images, resulting in an integrated signal within granules. The integrated signal was normalized to the values before rapamycin administration for quantification.

### Quantification and Statistical Analysis

Statistical parameters including the definition and exact values of N (number of cells and experiments), distribution and deviation are reported in figures and corresponding legends. Data are represented as mean ± SD using-tailed Welch’s t-tests. Statistical analyses were performed in Microsoft Excel.

---

**Supplementary Fig. 6.**
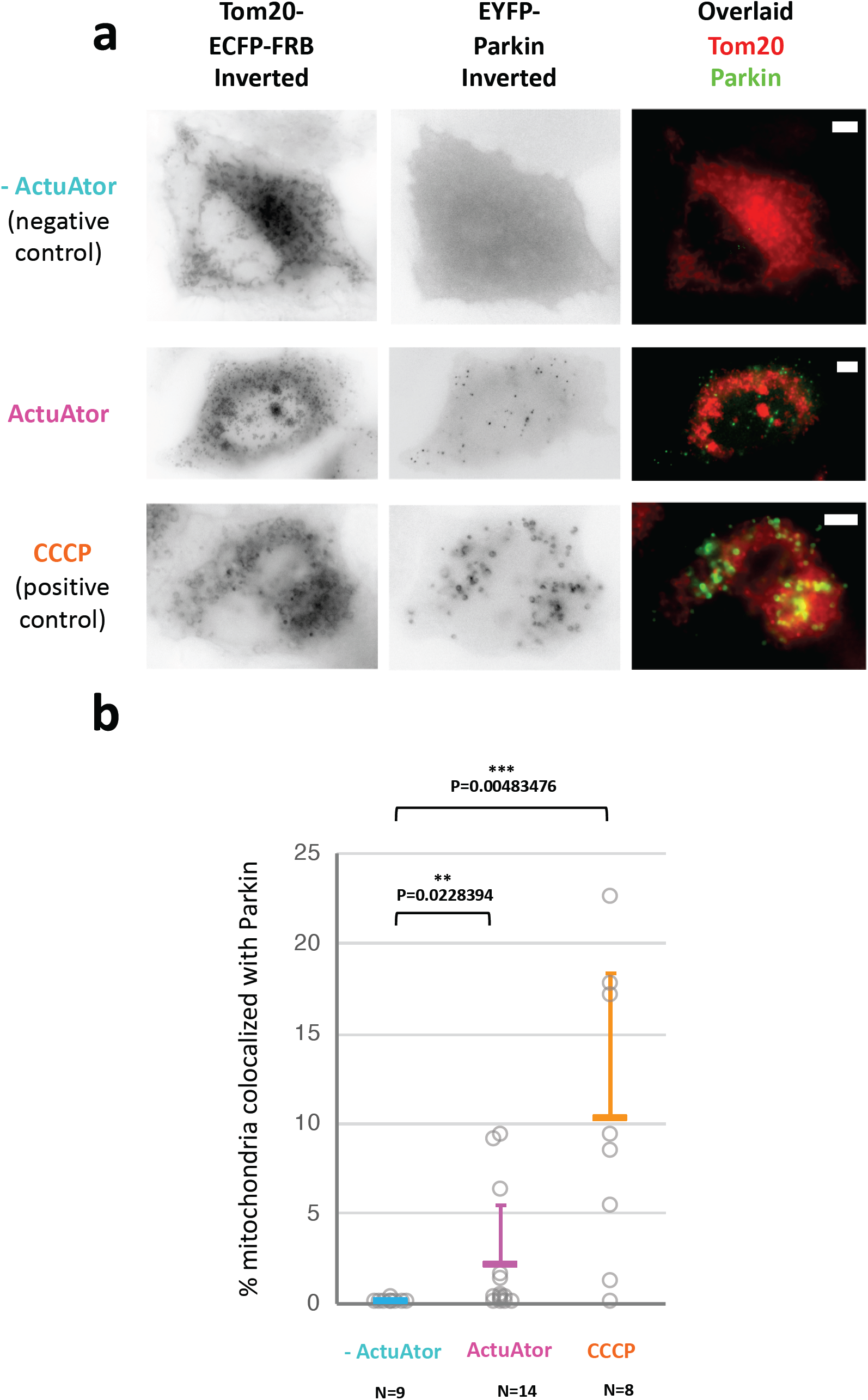

## References cited in Methods

Ermakova, Y.G., Bilan, D.S., Matlashov, M.E., Mishina, N.M., Markvicheva, K.N., Subach, O.M., Subach, F.V., Bogeski, I., Hoth, M., Enikolopov, G., et al. (2014). Red fluorescent genetically encoded indicator for intracellular hydrogen peroxide. Nat Commun 5, 5222.

Imamura, H., Nhat, K.P.H., Togawa, H., Saito, K., Iino, R., Kato-Yamada, Y., Nagai, T., and Noji, H. (2009). Visualization of ATP levels inside single living cells with fluorescence resonance energy transfer-based genetically encoded indicators. Proc. Natl. Acad. Sci. U.S.A. 106, 15651–15656.

Miyamoto, T., Rho, E., Sample, V., Akano, H., Magari, M., Ueno, T., Gorshkov, K., Chen, M., Tokumitsu, H., Zhang, J., et al. (2015). Compartmentalized AMPK signaling illuminated by genetically encoded molecular sensors and actuators. Cell Rep 11, 657–670.

Miyamoto, T., Rho, E., Kim, A., and Inoue, T. (2018). Cellular Application of Genetically Encoded Sensors and Impeders of AMPK. Methods Mol. Biol. 1732, 255–272.

Reitzer, L.J., Wice, B.M., and Kennell, D. (1979). Evidence that glutamine, not sugar, is the major energy source for cultured HeLa cells. J. Biol. Chem. 254, 2669–2676.

Rossignol, R., Gilkerson, R., Aggeler, R., Yamagata, K., Remington, S.J., and Capaldi, R.A. (2004). Energy substrate modulates mitochondrial structure and oxidative capacity in cancer cells. Cancer Res. 64, 985–993.

## REFERENCES

Banani, S.F., Lee, H.O., Hyman, A.A., and Rosen, M.K. (2017). Biomolecular condensates: organizers of cellular biochemistry. Nat. Rev. Mol. Cell Biol. 18, 285–298.

Beckerle, M.C. (1998). Spatial control of actin filament assembly: lessons from Listeria. Cell 95, 741–748.

van Bergeijk, P., Adrian, M., Hoogenraad, C.C., and Kapitein, L.C. (2015). Optogenetic control of organelle transport and positioning. Nature 518, 111–114.

Bi, E., and Zigmond, S.H. (1999). Actin polymerization: Where the WASP stings. Curr. Biol. CB 9, R160–163.

Boujemaa-Paterski, R., Gouin, E., Hansen, G., Samarin, S., Le Clainche, C., Didry, D., Dehoux, P., Cossart, P., Kocks, C., Carlier, M.F., et al. (2001). Listeria protein ActA mimics WASp family proteins: it activates filament barbed end branching by Arp2/3 complex. Biochemistry 40, 11390–11404.

Cameron, L.A., Footer, M.J., van Oudenaarden, A., and Theriot, J.A. (1999). Motility of ActA protein-coated microspheres driven by actin polymerization. Proc. Natl. Acad. Sci. U. S. A. 96, 4908–4913.

Castellano, F., Montcourrier, P., Guillemot, J.C., Gouin, E., Machesky, L., Cossart, P., and Chavrier, P. (1999). Inducible recruitment of Cdc42 or WASP to a cell-surface receptor triggers actin polymerization and filopodium formation. Curr. Biol. CB 9, 351–360.

Chakrabarti, R., Ji, W.-K., Stan, R.V., de Juan Sanz, J., Ryan, T.A., and Higgs, H.N. (2018). INF2-mediated actin polymerization at the ER stimulates mitochondrial calcium uptake, inner membrane constriction, and division. J. Cell Biol. 217, 251–268.

Chazotte, B. (2011). Labeling mitochondria with TMRM or TMRE. Cold Spring Harb. Protoc. 2011, 895–897.

Cramer, L.P., Mitchison, T.J., and Theriot, J.A. (1994). Actin-dependent motile forces and cell motility. Curr. Opin. Cell Biol. 6, 82–86.

De Vos, K.J., Allan, V.J., Grierson, A.J., and Sheetz, M.P. (2005). Mitochondrial function and actin regulate dynamin-related protein 1-dependent mitochondrial fission. Curr. Biol. CB 15, 678–683.

DeRose, R., Miyamoto, T., and Inoue, T. (2013). Manipulating signaling at will: chemically-inducible dimerization (CID) techniques resolve problems in cell biology. Pflüg. Arch. Eur. J. Physiol. 465, 409–417.

Diz-Muñoz, A., Fletcher, D.A., and Weiner, O.D. (2013). Use the force: membrane tension as an organizer of cell shape and motility. Trends Cell Biol. 23, 47–53.

Domann, E., Wehland, J., Rohde, M., Pistor, S., Hartl, M., Goebel, W., Leimeister-Wächter, M., Wuenscher, M., and Chakraborty, T. (1992). A novel bacterial virulence gene in Listeria monocytogenes required for host cell microfilament interaction with homology to the proline-rich region of vinculin. EMBO J. 11, 1981–1990.

Dufrêne, Y.F., Ando, T., Garcia, R., Alsteens, D., Martinez-Martin, D., Engel, A., Gerber, C., and Müller, D.J. (2017). Imaging modes of atomic force microscopy for application in molecular and cell biology. Nat. Nanotechnol. 12, 295–307.

Elosegui-Artola, A., Andreu, I., Beedle, A.E.M., Lezamiz, A., Uroz, M., Kosmalska, A.J., Oria, R., Kechagia, J.Z., Rico-Lastres, P., Le Roux, A.-L., et al. (2017). Force Triggers YAP Nuclear Entry by Regulating Transport across Nuclear Pores. Cell 171, 1397–1410.e14.

Elting, M.W., Suresh, P., and Dumont, S. (2018). The Spindle: Integrating Architecture and Mechanics across Scales. Trends Cell Biol. 28, 896–910.

Ermakova, Y.G., Bilan, D.S., Matlashov, M.E., Mishina, N.M., Markvicheva, K.N., Subach, O.M., Subach, F.V., Bogeski, I., Hoth, M., Enikolopov, G., et al. (2014). Red fluorescent genetically encoded indicator for intracellular hydrogen peroxide. Nat. Commun. 5, 5222.

Feng, Q., and Kornmann, B. (2018). Mechanical forces on cellular organelles. J. Cell Sci. 131.

Fradelizi, J., Noireaux, V., Plastino, J., Menichi, B., Louvard, D., Sykes, C., Golsteyn, R.M., and Friederich, E. (2001). ActA and human zyxin harbour Arp2/3-independent actin-polymerization activity. Nat. Cell Biol. 3, 699–707.

Friederich, E., Gouin, E., Hellio, R., Kocks, C., Cossart, P., and Louvard, D. (1995). Targeting of Listeria monocytogenes ActA protein to the plasma membrane as a tool to dissect both actin-based cell morphogenesis and ActA function. EMBO J. 14, 2731–2744.

Friedman, J.R., and Nunnari, J. (2014). Mitochondrial form and function. Nature 505, 335–343.

Golsteyn, R.M., Beckerle, M.C., Koay, T., and Friederich, E. (1997). Structural and functional similarities between the human cytoskeletal protein zyxin and the ActA protein of Listeria monocytogenes. J. Cell Sci. 110 (Pt 16), 1893–1906.

Gouin, E., Welch, M.D., and Cossart, P. (2005). Actin-based motility of intracellular pathogens. Curr. Opin. Microbiol. 8, 35–45.

Guet, D., Mandal, K., Pinot, M., Hoffmann, J., Abidine, Y., Sigaut, W., Bardin, S., Schauer, K., Goud, B., and Manneville, J.-B. (2014). Mechanical role of actin dynamics in the rheology of the Golgi complex and in Golgi-associated trafficking events. Curr. Biol. CB 24, 1700–1711.

Helle, S.C.J., Feng, Q., Aebersold, M.J., Hirt, L., Grüter, R.R., Vahid, A., Sirianni, A., Mostowy, S., Snedeker, J.G., Šarić, A., et al. (2017). Mechanical force induces mitochondrial fission. ELife 6.

Imamura, H., Nhat, K.P.H., Togawa, H., Saito, K., Iino, R., Kato-Yamada, Y., Nagai, T., and Noji, H. (2009). Visualization of ATP levels inside single living cells with fluorescence resonance energy transfer-based genetically encoded indicators. Proc. Natl. Acad. Sci. U. S. A. 106, 15651–15656.

Ingber, D.E. (2003). Mechanobiology and diseases of mechanotransduction. Ann. Med. 35, 564–577.

Isermann, P., and Lammerding, J. (2013). Nuclear mechanics and mechanotransduction in health and disease. Curr. Biol. CB 23, R1113–1121.

Iskratsch, T., Wolfenson, H., and Sheetz, M.P. (2014). Appreciating force and shape—the rise of mechanotransduction in cell biology. Nat. Rev. Mol. Cell Biol. 15, 825–833.

Itabashi, T., Takagi, J., Suzuki, K., and Ishiwata, S. (2013). Responses of chromosome segregation machinery to mechanical perturbations. Biophys. Nagoya-Shi Jpn. 9, 73–78.

Kassianidou, E., Kalita, J., and Lim, R.Y.H. (2019). The role of nucleocytoplasmic transport in mechanotransduction. Exp. Cell Res. 377, 86–93.

Kedersha, N., Ivanov, P., and Anderson, P. (2013). Stress granules and cell signaling: more than just a passing phase? Trends Biochem. Sci. 38, 494–506.

Kocks, C., Gouin, E., Tabouret, M., Berche, P., Ohayon, H., and Cossart, P. (1992). L. monocytogenes-induced actin assembly requires the actA gene product, a surface protein. Cell 68, 521–531.

Kocks, C., Hellio, R., Gounon, P., Ohayon, H., and Cossart, P. (1993). Polarized distribution of Listeria monocytogenes surface protein ActA at the site of directional actin assembly. J. Cell Sci. 105 (Pt 3), 699–710.

Komatsu, T., Kukelyansky, I., McCaffery, J.M., Ueno, T., Varela, L.C., and Inoue, T. (2010). Organelle-specific, rapid induction of molecular activities and membrane tethering. Nat. Methods 7, 206–208.

Kornmann, B. (2013). The molecular hug between the ER and the mitochondria. Curr. Opin. Cell Biol. 25, 443–448.

Korobova, F., Ramabhadran, V., and Higgs, H.N. (2013). An actin-dependent step in mitochondrial fission mediated by the ER-associated formin INF2. Science 339, 464–467.

Li, S., Xu, S., Roelofs, B.A., Boyman, L., Lederer, W.J., Sesaki, H., and Karbowski, M. (2015). Transient assembly of F-actin on the outer mitochondrial membrane contributes to mitochondrial fission. J. Cell Biol. 208, 109–123.

Liu, A.P. (2016). Biophysical Tools for Cellular and Subcellular Mechanical Actuation of Cell Signaling. Biophys. J. 111, 1112–1118.

Loisel, T.P., Boujemaa, R., Pantaloni, D., and Carlier, M.F. (1999). Reconstitution of actin-based motility of Listeria and Shigella using pure proteins. Nature 401, 613–616.

Luo, T., Mohan, K., Iglesias, P.A., and Robinson, D.N. (2013). Molecular mechanisms of cellular mechanosensing. Nat. Mater. 12, 1064–1071.

Mammoto, T., Mammoto, A., and Ingber, D.E. (2013). Mechanobiology and developmental control. Annu. Rev. Cell Dev. Biol. 29, 27–61.

Manor, U., Bartholomew, S., Golani, G., Christenson, E., Kozlov, M., Higgs, H., Spudich, J., and Lippincott-Schwartz, J. (2015). A mitochondria-anchored isoform of the actin-nucleating spire protein regulates mitochondrial division. ELife 4.

Mogilner, A., and Oster, G. (2003). Force generation by actin polymerization II: the elastic ratchet and tethered filaments. Biophys. J. 84, 1591–1605.

Moore, A.S., Wong, Y.C., Simpson, C.L., and Holzbaur, E.L.F. (2016). Dynamic actin cycling through mitochondrial subpopulations locally regulates the fission-fusion balance within mitochondrial networks. Nat. Commun. 7, 12886.

Nadappuram, B.P., Cadinu, P., Barik, A., Ainscough, A.J., Devine, M.J., Kang, M., Gonzalez-Garcia, J., Kittler, J.T., Willison, K.R., Vilar, R., et al. (2019). Nanoscale tweezers for single-cell biopsies. Nat. Nanotechnol. 14, 80–88.

Nakamura, H., Lee, A.A., Afshar, A.S., Watanabe, S., Rho, E., Razavi, S., Suarez, A., Lin, Y.-C., Tanigawa, M., Huang, B., et al. (2018). Intracellular production of hydrogels and synthetic RNA granules by multivalent molecular interactions. Nat. Mater. 17, 79–89.

Nikkhah, M., Edalat, F., Manoucheri, S., and Khademhosseini, A. (2012). Engineering microscale topographies to control the cell-substrate interface. Biomaterials 33, 5230–5246.

Norregaard, K., Metzler, R., Ritter, C.M., Berg-Sørensen, K., and Oddershede, L.B. (2017). Manipulation and Motion of Organelles and Single Molecules in Living Cells. Chem. Rev. 117, 4342–4375.

Pantaloni, D., Le Clainche, C., and Carlier, M.F. (2001). Mechanism of actin-based motility. Science 292, 1502–1506.

Pernas, L., and Scorrano, L. (2016). Mito-Morphosis: Mitochondrial Fusion, Fission, and Cristae Remodeling as Key Mediators of Cellular Function. In Annual Review of Physiology, Vol 78, D. Julius, ed. (Palo Alto: Annual Reviews), pp. 505-+.

Picard, M., Shirihai, O.S., Gentil, B.J., and Burelle, Y. (2013). Mitochondrial morphology transitions and functions: implications for retrograde signaling? Am. J. Physiol. Regul. Integr. Comp. Physiol. 304, R393–406.

Pickles, S., Vigie, P., and Youle, R.J. (2018). Mitophagy and Quality Control Mechanisms in Mitochondrial Maintenance. Curr. Biol. 28, R170–R185.

Pistor, S., Chakraborty, T., Niebuhr, K., Domann, E., and Wehland, J. (1994). The ActA protein of Listeria monocytogenes acts as a nucleator inducing reorganization of the actin cytoskeleton. EMBO J. 13, 758–763.

Protter, D.S.W., and Parker, R. (2016). Principles and Properties of Stress Granules. Trends Cell Biol. 26, 668–679.

Radoshevich, L., and Cossart, P. (2018). Listeria monocytogenes: towards a complete picture of its physiology and pathogenesis. Nat. Rev. Microbiol. 16, 32–46.

Rai, A.K., Rai, A., Ramaiya, A.J., Jha, R., and Mallik, R. (2013). Molecular Adaptations Allow Dynein to Generate Large Collective Forces inside Cells. Cell 152, 172–182.

Rowland, A.A., and Voeltz, G.K. (2012). Endoplasmic reticulum-mitochondria contacts: function of the junction. Nat. Rev. Mol. Cell Biol. 13, 607–625.

Sallee, N.A., Rivera, G.M., Dueber, J.E., Vasilescu, D., Mullins, R.D., Mayer, B.J., and Lim, W.A. (2008). The pathogen protein EspF(U) hijacks actin polymerization using mimicry and multivalency. Nature 454, 1005–1008.

Shivashankar, G.V. (2019). Mechanical regulation of genome architecture and cell-fate decisions. Curr. Opin. Cell Biol. 56, 115–121.

Skoble, J., Portnoy, D.A., and Welch, M.D. (2000). Three regions within ActA promote Arp2/3 complex-mediated actin nucleation and Listeria monocytogenes motility. J. Cell Biol. 150, 527–538.

Smith, G.A., and Portnoy, D.A. (1997). How the Listeria monocytogenes ActA protein converts actin polymerization into a motile force. Trends Microbiol. 5, 272–276.

Smith, G.A., Portnoy, D.A., and Theriot, J.A. (1995). Asymmetric distribution of the Listeria monocytogenes ActA protein is required and sufficient to direct actin-based motility. Mol. Microbiol. 17, 945–951.

Sniadecki, N.J. (2010). Minireview: A Tiny Touch: Activation of Cell Signaling Pathways with Magnetic Nanoparticles. Endocrinology 151, 451–457.

Suresh, S. (2007). Biomechanics and biophysics of cancer cells. Acta Biomater. 3, 413–438.

Tanase, M., Biais, N., and Sheetz, M. (2007). Magnetic tweezers in cell biology. In Cell Mechanics, Y.L. Wang, and D.E. Discher, eds. (San Diego: Elsevier Academic Press Inc), pp. 473–493.

Taslimi, A., Vrana, J.D., Chen, D., Borinskaya, S., Mayer, B.J., Kennedy, M.J., and Tucker, C.L. (2014). An optimized optogenetic clustering tool for probing protein interaction and function. Nat. Commun. 5, 4925.

Theriot, J.A., Mitchison, T.J., Tilney, L.G., and Portnoy, D.A. (1992). The rate of actin-based motility of intracellular Listeria monocytogenes equals the rate of actin polymerization. Nature 357, 257–260.

Várkuti, B.H., Képiró, M., Horváth, I.Á., Végner, L., Ráti, S., Zsigmond, Á., Hegyi, G., Lenkei, Z., Varga, M., and Málnási-Csizmadia, A. (2016). A highly soluble, non-phototoxic, non-fluorescent blebbistatin derivative. Sci. Rep. 6, 26141.

Vining, K.H., and Mooney, D.J. (2017). Mechanical forces direct stem cell behaviour in development and regeneration. Nat. Rev. Mol. Cell Biol. 18, 728–742.

Ward, M.W. (2010). Quantitative analysis of membrane potentials. Methods Mol. Biol. Clifton NJ 591, 335–351.

Weber, S.C., and Brangwynne, C.P. (2012). Getting RNA and protein in phase. Cell 149, 1188–1191.

Welch, M.D., Iwamatsu, A., and Mitchison, T.J. (1997). Actin polymerization is induced by Arp2/3 protein complex at the surface of Listeria monocytogenes. Nature 385, 265–269.

Welch, M.D., Rosenblatt, J., Skoble, J., Portnoy, D.A., and Mitchison, T.J. (1998). Interaction of human Arp2/3 complex and the Listeria monocytogenes ActA protein in actin filament nucleation. Science 281, 105–108.

Wheeler, R.J., and Hyman, A.A. (2018). Controlling compartmentalization by non-membrane-bound organelles. Philos. Trans. R. Soc. Lond. B. Biol. Sci. 373.

Yamada, T., Murata, D., Adachi, Y., Itoh, K., Kameoka, S., Igarashi, A., Kato, T., Araki, Y., Huganir, R.L., Dawson, T.M., et al. (2018). Mitochondrial Stasis Reveals p62-Mediated Ubiquitination in Parkin-Independent Mitophagy and Mitigates Nonalcoholic Fatty Liver Disease. Cell Metab. 28, 588–604.e5.

Youle, R.J., and van der Bliek, A.M. (2012). Mitochondrial Fission, Fusion, and Stress. Science 337, 1062–1065.

Zalevsky, J., Grigorova, I., and Mullins, R.D. (2001). Activation of the Arp2/3 complex by the Listeria acta protein. Acta binds two actin monomers and three subunits of the Arp2/3 complex. J. Biol. Chem. 276, 3468–3475.

Zhang, H., and Liu, K.-K. (2008). Optical tweezers for single cells. J. R. Soc. Interface 5, 671–690.

Zhang, K., Daigle, J.G., Cunningham, K.M., Coyne, A.N., Ruan, K., Grima, J.C., Bowen, K.E., Wadhwa, H., Yang, P., Rigo, F., et al. (2018). Stress Granule Assembly Disrupts Nucleocytoplasmic Transport. Cell 173, 958–971.e17.

