## Supplementary Information for "ActuAtor, a molecular tool for generating force in living cells: Controlled deformation of intracellular structures"

Hideki Nakamura, Elmer Rho, Daqi Deng, Shiva Razavi, Hideaki T. Matsubayashi, Takanari Inoue

### Contents:


- Supplementary Figures 1-6
- Figure Legends for Supplementary Figures 1-6
- Legends for Supplementary Movies 1-7

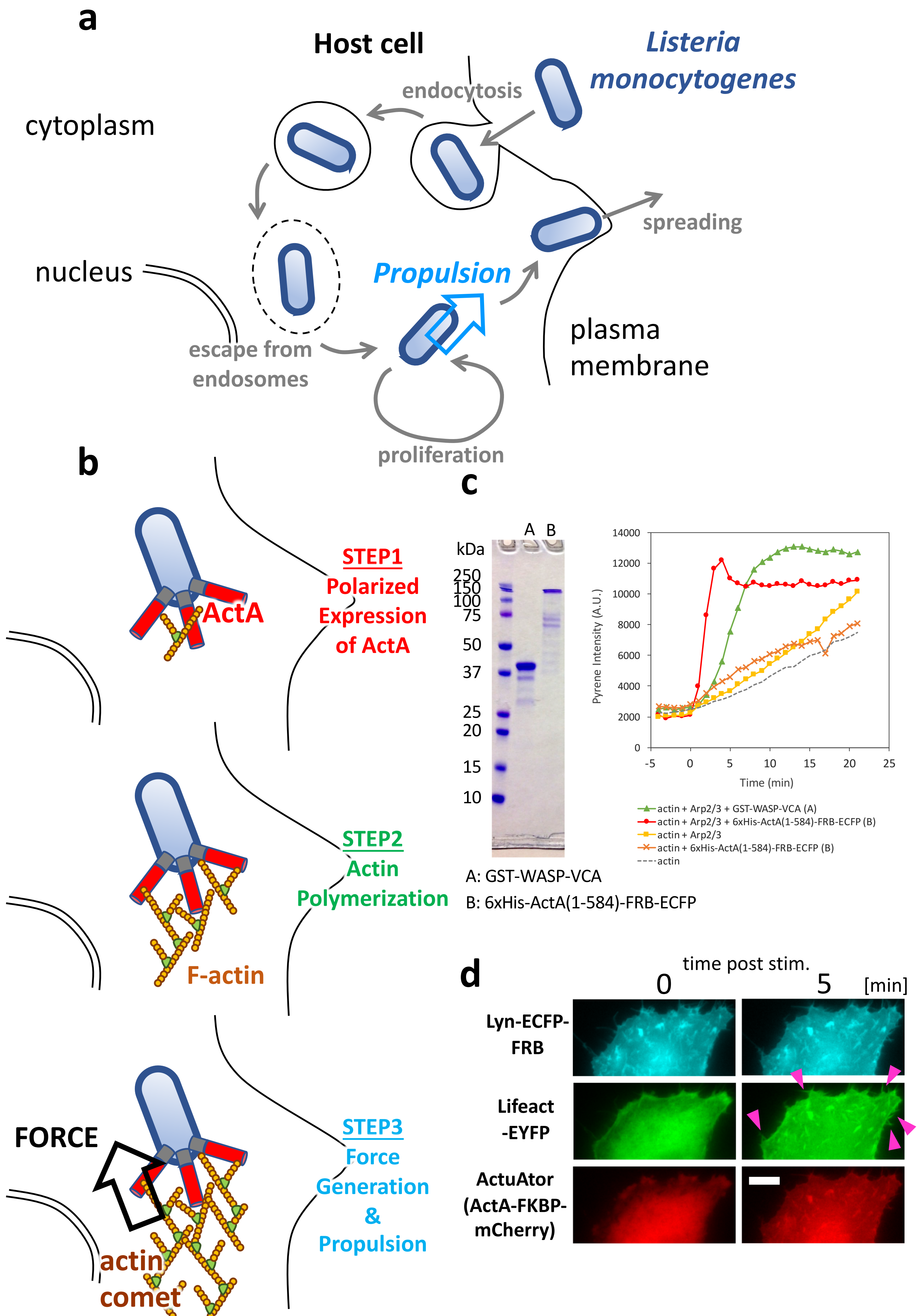


Supplementary Fig. 1 Nakamura et al

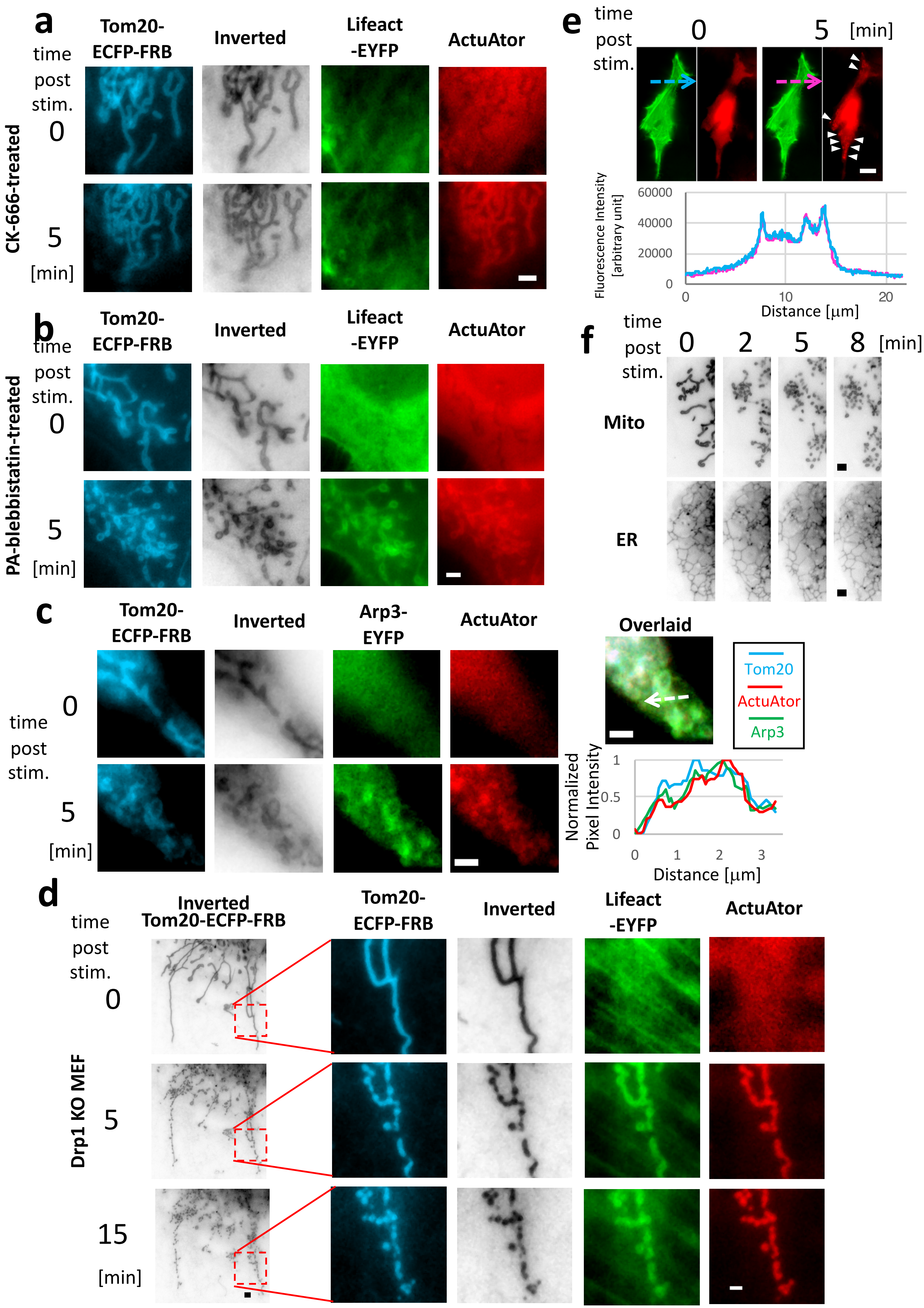


Supplementary Fig. 2 Nakamura et al

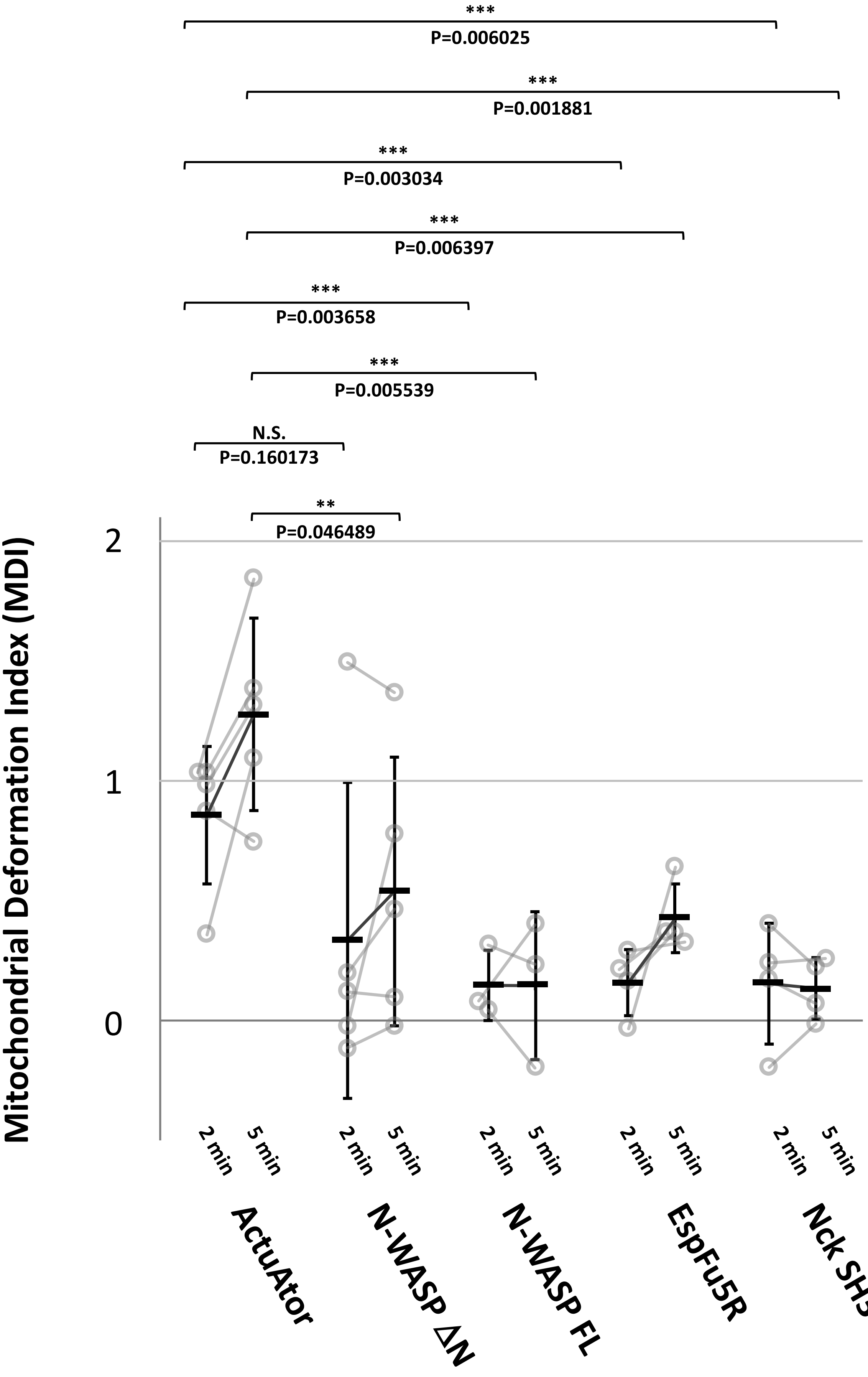


Supplementary Fig. 3 Nakamura et al

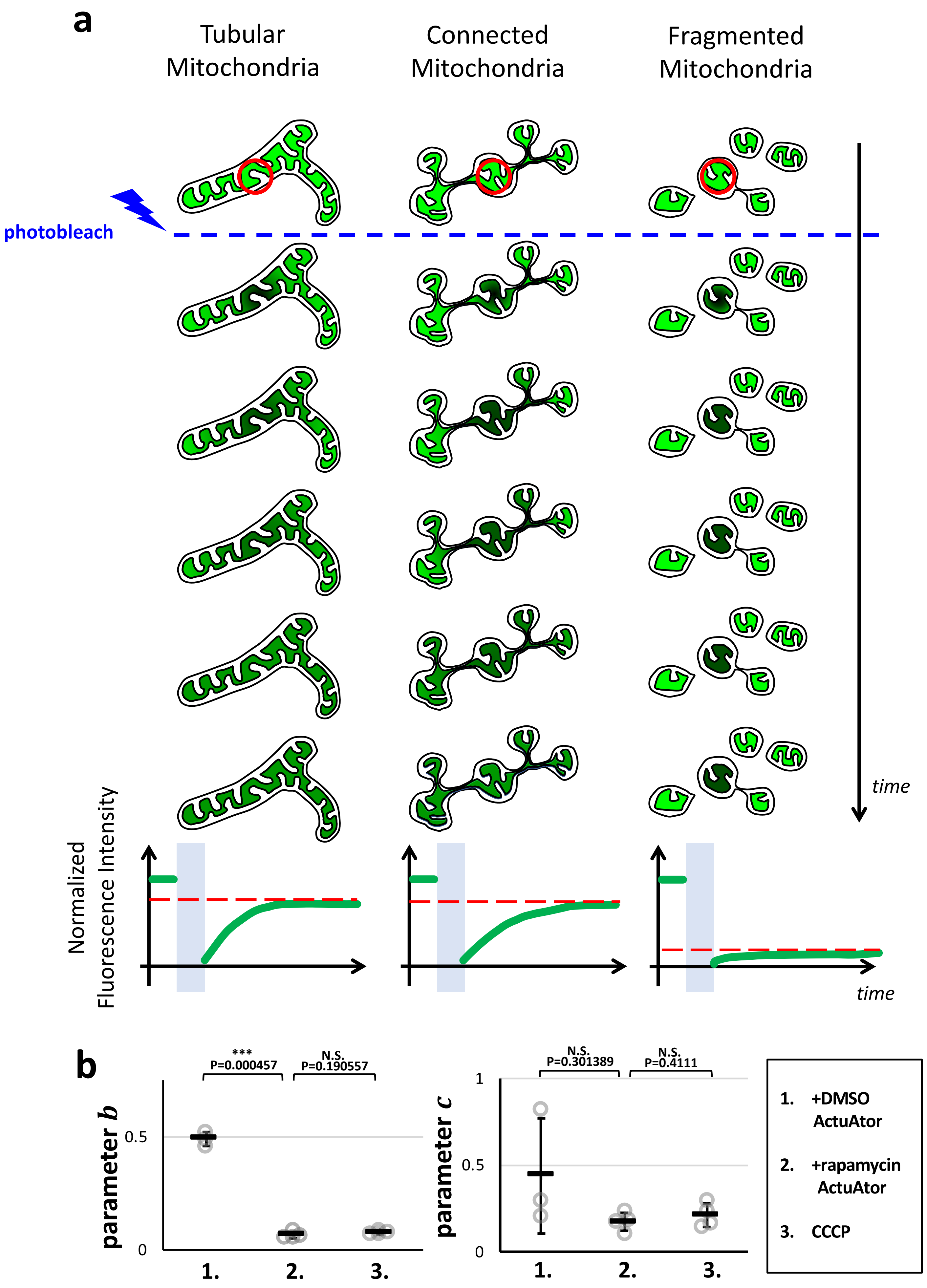


Supplementary Fig. 4 Nakamura et al

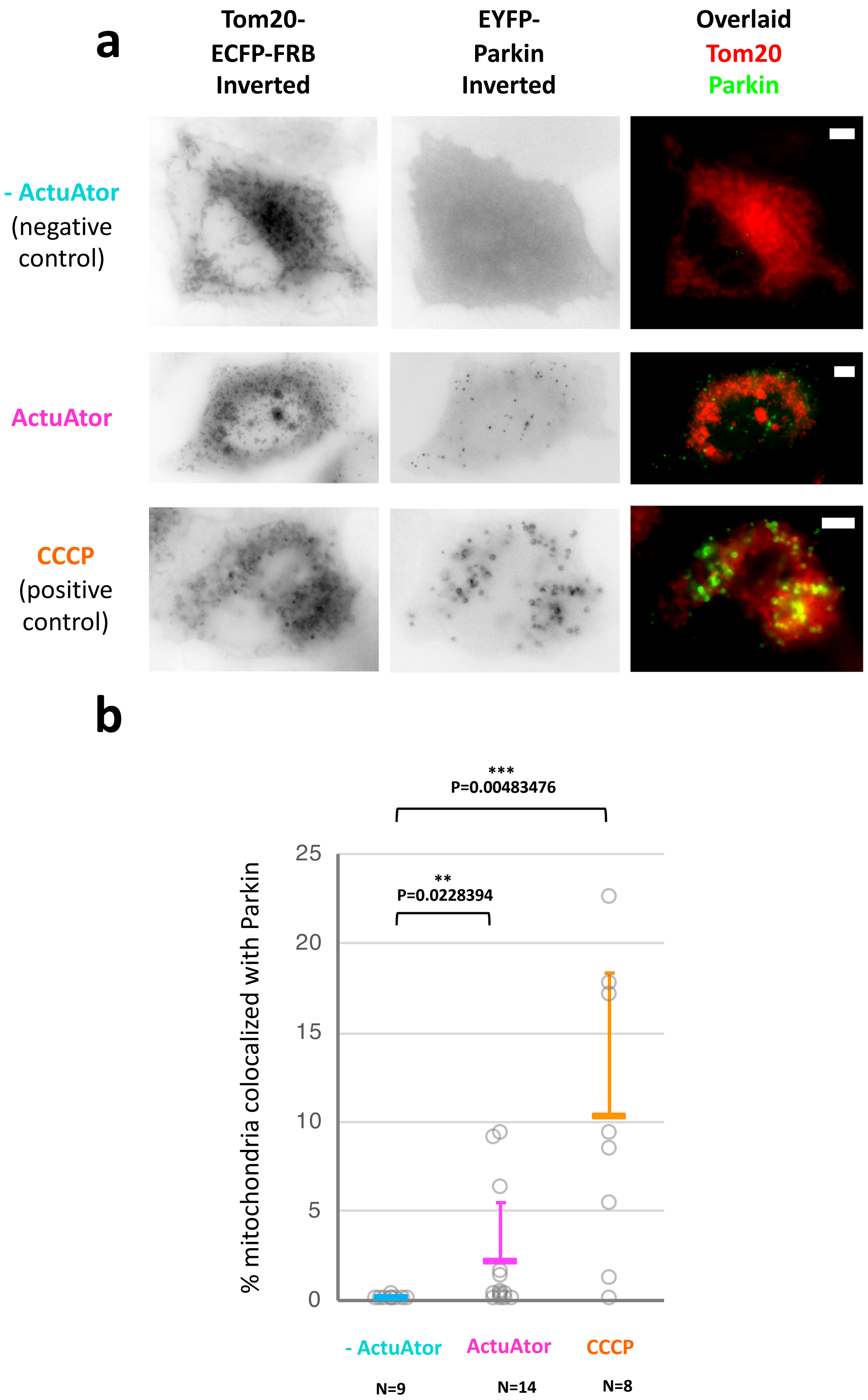


Supplementary Fig. 5 Nakamura et al

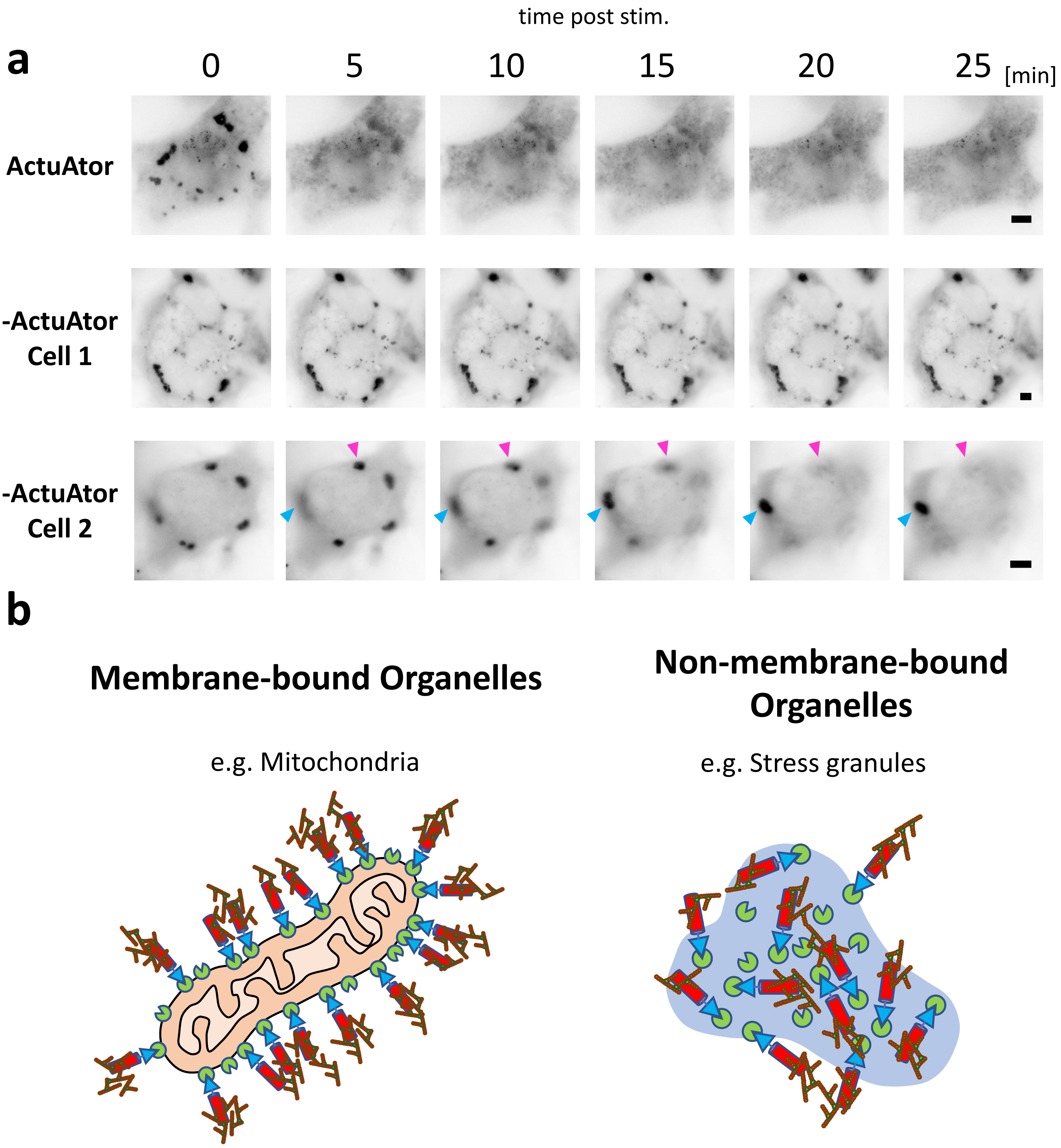


Supplementary Fig. 6 Nakamura et al

Legends for Supplementary Figures

**Supplementary Figure 1. Development of a novel tool, ActuAtor, that generates force in living cells based on a bacterial protein, ActA.**

a) Life cycle of *Listeria monocytogenes* in host cells. *Listeria* invades into the cytosol by endocytic entry into host cells followed by escape from endosomes. They proliferate in the cytosol and move around by hijacking actin polymerization of the host cell. The propulsion process is essential for the bacteria to escape from the host cell to spread across other cells in the tissue.

b) Mechanism of *Listeria* propulsion in the host cell cytosol. A bacterial membrane protein, ActA, is essential for the process. In the host cell cytosol, *Listeria* expresses ActA in a polarized manner (top panel). Extracellular domain of ActA then induces polymerization of host cell actin by functionally mimicking actin nucleation promoting factors of the host cell (middle panel). The polymerized actin polarization generates directional force exerted onto the bacteria, propelling them in the cytosol to realize bacterial motion (bottom panel).

c) Actin nucleation by the modified ActA peptide was confirmed *in vitro*. Consistent with previous *in vitro* studies of *Listeria* ActA, purified ActA(1-584) fused to FRB-ECFP showed Arp2/3 dependent actin polymerization activity in pyrene-actin assay. The kinetics was fast enough to indicate an overshooting peak, which is known to be observed when polymerization is more rapid than ATP hydrolysis and nucleotide exchange(Brooks and Carlsson, 2008).

d) ActuAtor-induced actin polymerization at plasma membrane led to formation of F-actin-rich microspikes. Fluorescence images of Lyn-ECFP-FRB (plasma membrane marker, cyan), Lifeact-EYFP (polymerized actin marker, green), and ActuAtor-FKBP-mCherry (ActuAtor peptide, red) right before (0 min) and five minutes (5 min) after rapamycin addition are shown. Upon stimulus, ActuAtor peptide was translocated to plasma membrane, where actin-rich microspikes are formed, as indicated by magenta arrowheads. Scale bar: 5 μm.

**Supplementary Figure 2. Mitochondria deformation by ActuAtor was dependent on Arp2/3, but not on myosin nor on mitochondria fission machinery.**

a) ActuAtor did not significantly deform mitochondria in HeLa cells pre-treated by Arp2/3 inhibitor, CK666. Mitochondria morphology (cyan and inverted monochrome panels), Lifeact signal (green) and ActuAtor peptide signal (red) right before (0 min) and five minutes after stimulus (5 min) are shown. Although ActuAtor was inducibly accumulated at mitochondria, increase in Lifeact signal was modest and no significant deformation was observed. Scale bar: 2 μm.

b) Deformation by ActuAtor in HeLa cells was not affected by myosin inhibitor, PA-blebbistatin. Scale bar: 2 μm.

c) Actin polymerization induced by ActuAtor involved Arp3. Arp3-EYP was accumulated on the surface of deformed mitochondria in HeLa cells, as shown in green panels. Scale bar: 2 μm.

d) Mitochondria deformation by ActuAtor did not require endogenous mitochohndria fission GTPase, Drp1. Mitochondria deformation in Drp1-KO MEF is shown. Scale bars: 2 μm.
e) ActuAtor did not significantly affect pre-existing actin cytoskeleton structures during mitochondria deformation. Fluorescence images of Lifeact-EYFP(green) and ActuAtor(red) in a HeLa cell right before (0 min) and five minutes after stimulus (5 min) are shown. Fluorescence intensity profiles along cyan and magenta broken-line arrows are plotted against distance in the lower panel in corresponding colors. Pre-existing F-actin-rich structures were not affected by ActuAtor recruitment, as demonstrated by the little change in profiles. Note that the focal plane was adjusted to clearly observe basal stress fibers, and mitochondrial accumulation as well as deformation are off-focus and only vaguely observable in ActuAtor channel (highlighted by white arrowheads). Scale bar: 10 μm.
f) Mitochondria deformation by ActuAtor did not severely affect the morphology of ER, which is closely associated with mitochondria. Inverted fluorescence intensity images of a mitochondria marker (Tom20-ECFP-FRB, Mito) and ER marker (EYFP-KDEL, ER) in a COS-7 cell at distinct time points are shown. ER tubular network remained mainly intact during drastic mitochondrial deformation induced by ActuAtor, although modest effects on ER morphology cannot be excluded. Scale bars: 2 μm.

**Supplementary Figure 3. ActuAtor outperformed previously reported actin nucleators in HeLa cells.** To quantify mitochondrial deformation, MDI (defined in Figure 1c) was calculated at two and five minutes after rapamycin addition for various actin nucleators including ActuAtor (ActuAtor, N=5 cells), N-terminally deleted N-WASP (N-WASP ΔN, N=5 cells), full-length N-WASP (N-WASP FL, N=3 cells), five tandem repeat of EspFu domain from a pathogenic *E. coli* strain (EspFu5R, N=4 cells), and SH3 domains from Nck (Nck SH3, N=4 cells). Deformation by Actuator was significantly more efficient than any other nucleators reported previously, although relatively modest deformation was observed with other nucleators. Statistical analysis was carried out by single-sided Welch's *t*-test. ***: $p<0.001$, **: $p<0.05$, N.S.: not significant.

**Supplementary Figure 4. Mitochondria connectivity evaluated by FRAP measurement.**
a) Schematic illustrations of FRAP measurements and their expected results in mitochondria with various level of connectivity. Of note, the actual size of connected space should be most reliably reflected in the asymptotic value after the total recovery (red broken line in the recovery plot in lower panels), which corresponds to the value of parameter $a$ in the formula in Figure 5c.
b) Fitted values for the two parameters, $b$ and $c$ in the formula in Figure 5c, are plotted. ***: $p<0.01$, N.S.: not significant.

**Supplementary Figure 5. Mitochondria deformation by ActuAtor affected mitophagy in the long term.**
a) Representative images of mitochondrial morphology (Tom20-ECFP-FRB, shown as inverted

images) and Parkin localization (EYFP-Parkin) in HeLa cells 24 hour after stimulus. Results without ActuAtor peptide recruitment (negative control, -ActuAtor,), those with ActuAtor (ActuAtor), and positive control experiment with CCCP treatment instead of rapamycin (CCCP) are shown. In the right column, overlaid images of mitochondria morphology (red) and Parkin speckles extracted by thresholding (green) are presented. Scale bars: 5 μm.

b) Colocalization of a mitophagy marker, Parkin, with mitochondria quantified as percentage of mitochondrial area colocalized with Parkin speckles. Individual data is plotted as a grey circular marker, while mean + standard deviation is shown as marker and an error bar in relevant colors. N=9, 14, and 8 cells for -ActuAtor, ActuAtor, and CCCP conditions, respectively. Mitochondria deformed by ActuAtor tended to colocalize modestly but significantly more frequently with Parkin.

**Supplementary Figure 6. Stress granule dispersion by ActuAtor.**

a) Recruitment of ActuAtor peptide or negative control peptide lacking ActuAtor to stress granules by TIA1. Independent marker, EYFP-PABP1, was used to monitor stress granule morphology. Inverted fluorescence images of the marker at distinct time points are shown. Although the dispersion by ActuAtor was robust (ActuAtor), negative control peptide accumulation on the mitochondria led to diverse phenotypes (-ActuAtor Cell1, -ActuAtor Cell2), as reflected in large deviation in the plot in Figure 7c. In one cell, the recruitment caused significant dispersion of the majority of the granules (-ActuAtor Cell2), leading to disappearance of granules according to the quantification in Figure 7c, while no significant change was observed in another cell (-ActuAtor Cell1). Interestingly, the dispersing effect may not be uniform even across granules in a single cell, implied by the coexistence of dispersing granule (magenta arrowhead) and emerging or growing granule (cyan arrowhead) in the same cell. Scale bars: 5 μm.

b) Schematic illustrations for ActuAtor-induced deformation of conventional organelles (left panel) and dispersion of non-membrane-bound organelle such as stress granules (right panel). Unlike the case of conventional organelles, induced actin polymerization during stress granule dispersion takes place all across the granules.

### Supplementary Movies

#### Supplementary movie S1. Mitochondria deformation by ActuAtor

Tom20-ECFP-FRB(mitochondria morphology marker, cyan), Lifeact-EYFP(F-actin marker, green), and ActuAtor(red) images from the same data as Figure 1b are shown as a movie.

#### Supplementary movie S2. Mitochondria deformation across an entire cell by ActuAtor

Inverted images of Tom20-ECFP-FRB(mitochondria morphology marker) across a representative COS-7 cell are shown as a movie.

#### Supplementary movie S3. Golgi apparatus deformation by ActuAtor

ECFP-FRB-giantin(Golgi apparatus marker, cyan), Lifeact-EYFP(F-actin marker, green), and ActuAtor(red) are shown as a movie. The data is the same as the one shown in Figure 3.

#### Supplementary movie S4. Actin-rich patch in a nucleus deformed by ActuAtor

ECFP-FRB-Sec61B(endoplasmic reticulum and outer nuclear membrane marker, cyan), Lifeact-EYFP(F-actin marker, green), and ActuAtor(red) are shown as a movie. The data is the same as the one shown in Figure 4a.

#### Supplementary movie S5. Actin-rich spikes found in cells with nuclear rupture induced by ActuAtor

ECFP-FRB-Sec61B(endoplasmic reticulum and outer nuclear membrane marker, cyan), Lifeact-EYFP(F-actin marker, green), ActuAtor(red), and iRFP713-NLS(soluble nucleus marker, magenda) are shown as a movie. The data is the same as the one shown in Figure 4b and Supplementary Figure 4.

#### Supplementary movie S6. Actin-rich finger-like protrusion found in nucleus deformed by ActuAtor

ECFP-FRB-Sec61B(endoplasmic reticulum and outer nuclear membrane marker, cyan), Lifeact-EYFP(F-actin marker, green), and ActuAtor(red) are shown as a movie. The data is the same as the one shown in Figure 4c.

#### Supplementary movie S7. Stress granule dispersion induced by ActuAtor

ECFP-FRB-TIA1(stress granule marker, cyan), Lifeact-EYFP(F-actin marker, green), and ActuAtor(red) are shown as a movie. The data is the same as the one shown in Figure 7a.
